# *A counter-enzyme complex regulates* glutamate metabolism in *Bacillus subtilis*

**DOI:** 10.1101/2021.04.12.439528

**Authors:** Vijay Jayaraman, D. John Lee, Nadav Elad, Shay Vimer, Michal Sharon, James S. Fraser, Dan S. Tawfik

**Author notes:** Corresponding author(s) e-mail address (es). These authors contributed equally (D. John Lee, Nadav Elad).

## Abstract

Multi-enzyme assemblies composed of metabolic enzymes catalyzing sequential reactions are being increasingly studied. Here, we report the discovery of a 1.6 megadalton multi-enzyme complex from *Bacillus subtilis* composed of two enzymes catalyzing opposite rather than sequential reactions (“counter-enzymes”): glutamate synthase (GltAB), and glutamate dehydrogenase (GudB), that make and break glutamate, respectively. *In vivo* and *in vitro* studies show that the primary role of complex formation is to inhibit GudB’s activity as this enzyme is constitutively expressed including in glutamate-limiting conditions. Using cryo-electron microscopy, we elucidated the structure of the complex and the basis of GudB’s inhibition. Finally, we show that this complex that exhibits unusual oscillatory progress curves is a necessity for planktonic growth in glutamate-limiting conditions, but is also essential for biofilm growth in glutamate-rich media, suggesting a regulatory role at fluctuating glutamate concentrations.

## Introduction

Metabolites are key cellular resources and their homeostasis is primarily achieved by regulating enzyme activities at various levels: transcription, translation, and/or the protein level. While regulation of transcription and translation prevents the waste of resources and generally enables near-complete silencing of enzyme activity, regulation at the protein level enables rapid changes in metabolite levels. Regulatory modes at these various levels are crucial for the fitness of the organism (Chubukov et al., 2013; Curi et al., 2016; Metallo and Vander Heiden, 2013). At the protein level, enzymes are typically regulated by small molecules binding to an allosteric site, and/or by post-translational modification. Evidence is accumulating, however, of transient enzyme complexes, or metabolons, that allow high metabolic efficiency through clustering of enzymes and compartmentalization of reaction intermediates, and may also enable tighter control of enzyme activity (Zhang and Fernie, 2020). In this study, we describe a unique mode of regulation where two central metabolic enzymes that catalyze opposite reactions (“counter-enzymes”) form a transient complex under glutamate-poor conditions. This mode of regulation which provides a robust, ultra-sensitive response has so far been seen only in signaling pathways (Hart and Alon, 2013).

Our study arose from the search for the regulatory mechanism of glutamate dehydrogenase, GudB, from *B. subtilis*. Glutamate dehydrogenases catalyze the interconversion of glutamate to α-ketoglutarate (AKG) and ammonia, hence reside at the crossroads of carbon and nitrogen metabolism. Glutamate, is the major nitrogen reservoir and can provide up to 85% of the cellular nitrogen requirement (Lin et al., 1996). Moreover, glutamate has multiple other roles: intracellular pH buffering, maintenance of osmolarity, and functions as counter ion to potassium (**Figure 1A**) (Yan et al., 1996; Young and Ajami, 2000). Glutamate (Glu) is also one of the most abundant metabolites with cellular concentrations up to 150 mM (Bennett et al., 2009; Van Eunen et al., 2010; Tempest et al., 1970). In contrast, AKG is maintained at low concentrations, in the range of a few millimolars, and has rapid turnover (Huergo and Dixon, 2015). In most Bacilli species, the Glu/AKG balance is maintained by the activity of catabolic, NAD^+^-dependent, dehydrogenases that degrade glutamate, with GudB being the key player in *B. subtilis* (Noda-Garcia et al., 2017). The other enzyme that directly affects the Glu/AKG balance is glutamate synthase (GltAB), a heterodimeric enzyme composed of two subunits GltA and GltB, which acts in the opposite direction to GudB. Besides these two enzymes, transaminases that are coupled to the Glu/AKG exchange also affect Glu/AKG balance (Fisher and Magasanik, 1984; Hu et al., 1999; Whatmore et al., 1990) (**Figure 1A**).

**Figure 1.**
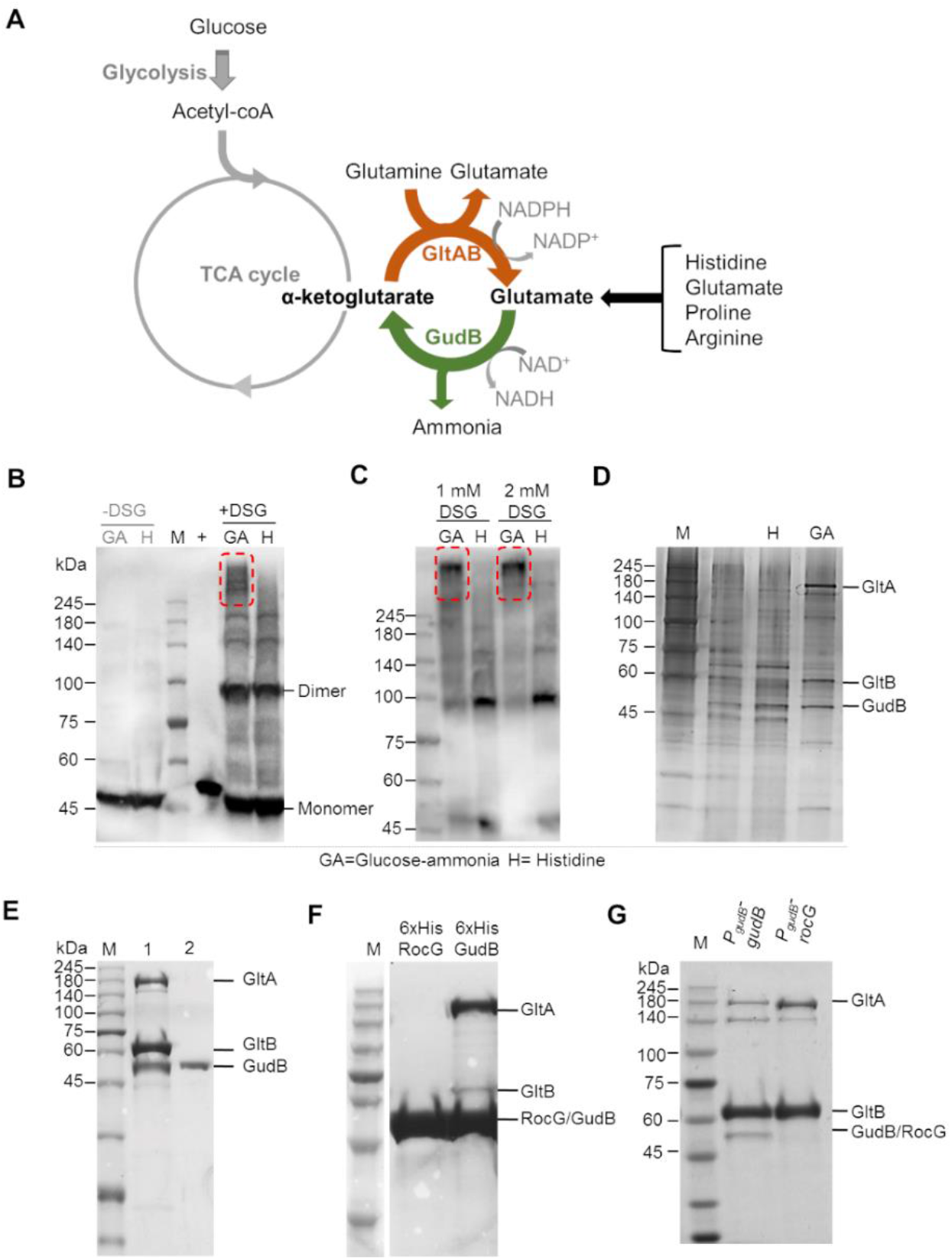
GudB interacts with GltAB in glutamate poor growth conditions. **A)** Schematic showing the key reactions involved in glutamate metabolism in *Bacillus subtilis*. Amino acids like proline, arginine, and histidine when provided as the sole C/N source are catabolized via glutamate. In contrast, growth on glucose as C source demands glutamate synthesis (via AKG). **B)** Western blot using anti-GudB antibodies indicating similar expression levels of GudB in *B. subtilis* cells grown in glucose-ammonia (GA) and Histidine (H) (-DSG). Upon treating with a chemical crosslinker (DSG, 0.5 mM), high molecular weight species that include GudB can be seen in cells grown on glucose-ammonia (GA, highlighted in red frame) but not on histidine (H). Recombinant GudB served as a positive control (+). **C)** The high molecular weight species of GudB are clearly seen in Western analysis of lysates from cells grown on glucose-ammonia (as in C) yet treated with higher concentration of DSG (1mM/2mM). **D)** Immunoprecipitation of GudB indicated co-elution of GltA and GltB in glucose-ammonia but not in histidine. The eluates from the pulldown was subjected to SDS-PAGE and stained with silver nitrate. **E)** SDS-PAGE showing co-elution of GudB and GltA upon purification of Strep-GltB from *B. subtilis* cells grown in glucose-ammonia (Lane 1; Lane 2 shows purified recombinant Strep tagged-GudB). **F)** Co-purification of GltA and GltB upon pulldown of His-tagged GudB but RocG. **G)** GudB but not RocG co-eluted upon pulldown of Strep-GltB from strains expressing either GudB or RocG from the constitutive *gudB* promoter (Gpt).

While not much is known about the post-translational regulation of glutamate synthases, glutamate dehydrogenases are among the most tightly regulated enzymes – they are renowned, if not infamous, for their complex allosteric regulatory features (Engel, 2011). The mammalian enzymes are regulated by nucleotides (Li et al., 2012), while some bacterial enzymes are regulated by amino acids such as leucine (Tomita et al., 2011). The *B. subtilis* GudB is constitutively expressed (Belitsky and Sonenshein, 1998; Noda-Garcia et al., 2017), and at high levels, and hence regulation at the protein level is critical. However, nucleotides, amino acids, and other key metabolites we have tested *in vitro* have all failed to regulate GudB’s activity. Instead, we observed that, *in vitro*, GudB equilibrates between an inactive dimer and an active hexamer in a glutamate- and pH-dependent manner. The hexamer’s instability suggested that its dissociation is the key to suppression of GudB’s activity when glutamate must be synthesized rather than catabolized (Noda-Garcia et al., 2017).

Aiming to test this hypothesis, we cross-linked GudB in *B. subtilis* cells grown in different media thus probing its oligomeric state at conditions where glutamate is either catabolized or synthesized. To our surprise, the trend was opposite of the expected one – high-M.W. species, that by our proposed model would indicate active hexamers, occurred under conditions where GudB should remain silent. Further investigations led to an even more surprising result: GudB is silenced by interacting with its “counter-enzyme”, glutamate synthase, GltAB. We further found that this complex occurs conditionally, at low glutamate when GltAB is expressed. Enzyme kinetics, native mass spectrometry, and high-resolution cryo-EM structures, revealed the molecular basis of this unusually large (1.6 MDa) and unique regulatory protein complex that exhibits oscillatory kinetics. Using genetic modifications, including the expression of an inactive GltAB variant, we show that, as expected, under planktonic growth complex formation is essential primarily in glutamate-poor conditions. However, we also found that this complex is needed for optimal biofilm development even in a glutamate-rich medium, suggesting that this counter-enzymes assembly may mediate the previously observed spatiotemporal glutamate catabolism in biofilms (Liu et al., 2015).

## Results

### GudB interacts with GltAB in glutamate-poor growth conditions

*B. subtilis* grows well in both glutamate-rich and glutamate-poor media. The former occurs when the carbon-nitrogen source is glutamate, or other amino acids like histidine, proline, or arginine that are catabolized via glutamate as intermediate (**Figure 1A**) – in our experiments, histidine mostly represented the glutamate-rich condition. The glutamate-poor condition is exemplified by growth on glucose-ammonia, a condition where glutamate must be synthesized. Oddly, the enzyme levels of GudB, the primary catabolic glutamate dehydrogenase in *B. subtilis*, are the same in both histidine and glucose-ammonia (**Figure 1B**, -DSG). Based on previous *in vitro* studies on GudB (Noda-Garcia et al., 2017), we anticipated, however, that in histidine, the enzyme would assemble into hexamers (the active state) while in glucose-ammonia it would dissociate to monomers/dimers (the inactive state).

To probe the *in vivo* oligomeric state of GudB, we performed Western blotting using anti-GudB antibody of cells grown in glutamate poor/rich conditions that were treated with a membrane permeable crosslinker (**Figure 1B**, +DSG). Surprisingly, in glucose-ammonia we observed bands with molecular weight similar, or possibly greater than expected for hexameric GudB (∼280 kDa), while in histidine the bands indicated dissociated monomer/dimer species. At higher cross linker concentration (**Figure 1C**), this discrepancy is even more evident: in cells grown in glucose-ammonia almost all of GudB was present as high-MW species. This species is specific to glutamate-poor media like glucose-ammonia, and not seen in any glutamate-rich medium, such as glutamate, histidine, arginine, or glycerol-glutamate (**Figure S1, A**).

In principle, the high-MW species observed in glucose-ammonia could be higher order oligomeric states of GudB or a complex of GudB with other proteins. To differentiate between the two possibilities, we performed antibody-based pulldowns of GudB from lysates of cells grown in either glutamate-rich or glutamate-poor conditions. Eluates from pulldowns contained at least two proteins in addition to GudB, with molecular weight around ∼160 kDa and ∼60 kDa, that were not observed in cell grown in the glutamate-rich medium (**Figure 1D**). Shotgun proteomics of the eluates indicated that these proteins are GltA (168.7 kDa) and GltB (54.8 kDa). Both proteins were significantly enriched in eluates obtained from glucose-ammonia grown cells compared to histidine grown ones (≤167 fold with crosslinker, and ≤ 21-fold without; **Table S1**). Finally, expression and pulldown of TwinStrep-tagged GltB (GltB-TS) from cells grown on glucose-ammonia also indicated the formation of a ternary GudB-GltAB complex (**Figure 1E**).

Why is the GudB-GltAB complex observed upon growth in glucose-ammonia but not in histidine? Unlike GudB, GltAB expression is highly regulated (Picossi et al., 2007; Smaldone et al., 2012). Indeed, GltAB is exclusively expressed in glutamate-poor conditions as shown by Western blots using strep tag antibodies against Strep-GltB (**Figure S1B**). Thus, it seems that in glutamate-rich conditions the GudB-GltAB complex does not form simply because GltAB is not present.

### The interaction with GltAB is paralog specific

GudB is not the only catabolic glutamate dehydrogenase in *B. subtilis*. Its closely related paralog, RocG, shares 75 % sequence identity and an effectively identical hexameric structure (Gunka et al., 2010). However, RocG is expressed at low levels (up to 30-fold lower compared to GudB), and its transcription is tightly regulated (triggered by arginine and removal of catabolic repression). Further, placement of RocG under *gudB*’s promoter indicated that RocG lacks the post-translational regulatory mode exhibited by GudB (Noda-Garcia et al., 2017). However, what this mode was remained unclear. We therefore compared the ability of RocG and GudB to bind GltAB by reciprocal pulldowns. *In vitro*, we observed that while GltAB co-elutes with GudB, no such interaction is observed with RocG (**Figure 1F**). To check if RocG could interact with GltAB *in vivo*, we used strains where either GudB or RocG were expressed from *gudB*’s constitutive promoter (namely, the wild-type *gudB* gene, and a replacement of the original ORF by *rocG*, respectively). Pulldowns of GltB from these strains reaffirmed that GltAB binds GudB but not RocG (**Figure 1G**). This result is in agreement with the promoter-ORF swaps of these paralogous genes (RocG protein under *gudB* promoter, and the reverse) showing severe growth defects (Noda-Garcia et al., 2017). This shows that in the absence of regulation at the level of gene/protein expression, GudB might be regulated by GltAB binding. Further, given the lack of interaction of RocG with GltAB, strains consitutively expressing RocG were used as a control in the growth phenotype experiments described below.

### GltAB negatively regulates GudB activity *in vivo*

Having established that GudB forms a complex with its counter-enzyme, GltAB, we next aimed to decipher the *in vivo* relevance of GudB-GltAB complex formation. Growth profiling of the individual knockout strains, Δ*gudB*, Δ*gltA*, and Δ*gltB*, in different media indicated non–overlapping requirements for the enzymatic activity of GudB and GltAB, *i.e.*, under conditions where GltAB is essential, GudB is dispensable (**Figure 2A**) and vice-versa (**Figure 2B,C**; note that removal of either of GltAB’s subunits leads to complete loss of the synthase activity). Specifically, the catabolic GudB is essential when glutamate (**Figure 2B**), or other amino acids such as histidine (**Figure 2C**) that are catabolized via glutamate, are present as the sole carbon source. Conversely, the anabolic GltAB is essential when glutamate needs to be synthesized de novo, as indicated by the inability of either Δ*gltA* or Δ*gltB* to grow on glucose-ammonia (**Figure 2B**). Finally, neither GudB nor GltA/B are required when glucose-ammonia is supplemented with glutamate (**Figure 2D**). Given the orthogonal functions, we surmised that binding of GltAB could be inhibiting GudB’s activity to avoid the futile cycle of glutamate synthesis and breakdown. If this is indeed the case, then GudB-GltAB complex formation should be deleterious under conditions where glutamate is the sole carbon source and GudB’s activity is pivotal.

**Figure 2:**
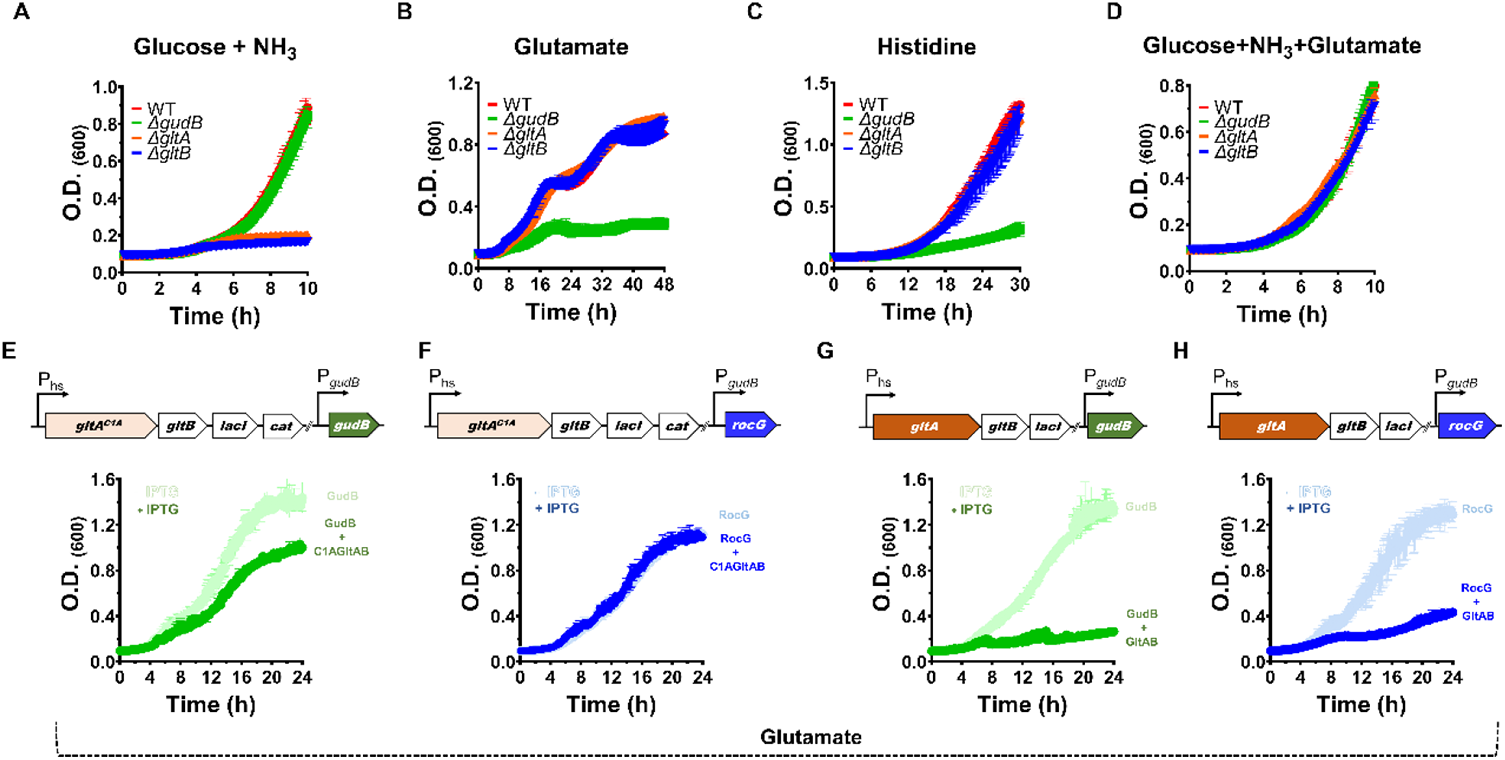
The phenotypic effects of GltAB and GudB and their interaction. **(A-D)** Growth profiling of the denoted knockout strains indicate that GltAB’s glutamate dehydrogenase activity is essential for growth in glucose-ammonia (panel A) while GudB’s glutamate dehydrogenase activity is essential in glutamate (panel B) and histidine (panel C). Under conditions where both glucose and glutamate are available neither of these two activities is essential (panel D). **(E-H)** Growth inhibition in glutamate medium upon expression of wild-type GltAB, and its inactive mutant (GltA^C1A^), from an IPTG inducible hyper-spank promoter. The scheme above each panel shows the genotype of the corresponding strain. Expression of GltAB^C1A^ causes growth suppression in a strain co-expressing GudB (panel E) but not in a strain co-expressing RocG (panel F). These effects are in agreement with GltA interacting with GudB but not with RocG (Figure 1F and 1G). In contrast, expression of functional GltA causes growth suppression in both the strains (GudB, panel G and RocG, panel H) likely due to futile cycling of making and breaking glutamate. Error bars in all panels represent SD from three independent growth experiments.

To test the above hypothesis, GltAB was placed under an inducible promoter, allowing it to be expressed also when glutamate is present in the medium. However, a growth defect observed in this case could also be because of futile cycling of glutamate instead of, or in addition to, GudB’s silencing. To circumvent the latter, we generated a strain, *P_hs_*-*gltAB*^C1A^, wherein GudB is expressed from its endogenous constitutive promoter and an inactive mutant of GltAB (GltAB^C1A^) is expressed from an IPTG-inducible hyperspank promoter. In this mutant (GltAB^C1A^), the N-terminal cysteine residue that catalyzes GltA’s amidotransferase activity (hydrolysis of glutamine to release ammonia that in turn reacts with AKG to generate glutamate) is mutated to alanine, and accordingly this mutant is devoid of glutamate synthase activity (**Figure S1 C**). This mutant does, however, bind GudB (**Figure S1 D**). IPTG-dependent expression of GltAB was verified in the *P_hs_*-*gltAB*+*gudB* strain, by showing that this strain grows in glucose-ammonia only in the presence of IPTG (**Figure S2**). These inducible GltAB genes (both Wt and C1A mutant) were also introduced in a strain expressing RocG under the *gudB* promoter (*P_hs_-gltAB*+*PgudB-rocG* and *P_hs_-gltAB*^C1A^+*PgudB-rocG*). The latter served as negative control because as elaborated in the above section, RocG does not interact with GltAB (**Figure 1F, 1H**). The growth profile of these strains, when grown on glutamate as the sole carbon and nitrogen source, with or without transgene induction, are presented in **Figure 2E-H**.

The key observation from this experiment is the growth defect seen in the strain, *P_hs_-gltAB*^C1A^+*gudB*, expressing GltAB^C1A^ alongside GudB, in a glutamate-based medium (**Figure 2E**). Here, it is the suppression of GudB’s activity by GltAB that inhibits growth, as validated by pulldowns of the GudB-GltAB^C1A^ complex under the same growth condition (**Figure S1D**). Further, no growth inhibition was observed when the inactive GltAB^C1A^ was co-expressed with RocG (**Figure 2F**; and accordingly, GltAB^C1A^ was pulled-down on its own, **Figure S1D**). As expected, a severe growth defect is observed in the strain expressing GudB and a functional GltAB (**Figure 2G**). However, a similar defect is observed with RocG, which does not interact with GltAB (**Figure 2H**) suggesting that growth inhibition in this case is mostly the result of futile cycling, *i.e.*, of making and breaking glutamate when a dehydrogenase and synthase are both present and functional (hence “burning” NADPH, and ATP due to glutamine hydrolysis). Taken together, these results support a key regulatory role for the GudB-GltAB complex, and specifically, in silencing GudB under glutamate-poor conditions, *i.e.*, when glutamate synthesis is essential and its breakdown is deleterious. Next, we turned to *in vitro* studies, aiming to examine the effect of complex formation on the activity of GudB and GltAB.

### GltAB binding alters the kinetic properties of GudB

To understand the kinetic effects of complex formation, the complex, and the individual enzymes, were purified from *B. subtilis* (**Figure 3A**) and used for steady-state kinetic measurements. As expected, the complex exhibited either glutamate synthase or dehydrogenase activity depending on which substrates were added (**Figure 3B**), and the steady state kinetic parameters were accordingly derived from initial velocity measurements (**Table 3C**; **Figure S3 A-F**). The activity of GltAB is essentially the same on its own and in complex with GudB. However, GudB showed altered behaviour: the KM value for glutamate is ∼19 times higher in the complex (7.3 mM for free GudB, **Figure 3C**, versus 138 mM for the GltAB bound form, **Figure 3D**). Further, while free GudB showed negative cooperativity with respect to glutamate as the substrate, the GltAB-bound form showed weak yet reproducible positive cooperativity (H=1.3; **Figure 3D**). Both these results support the notion that GltAB binding suppresses GudB’s activity, in accordance with the in vivo results (the section above).

**Figure 3.**
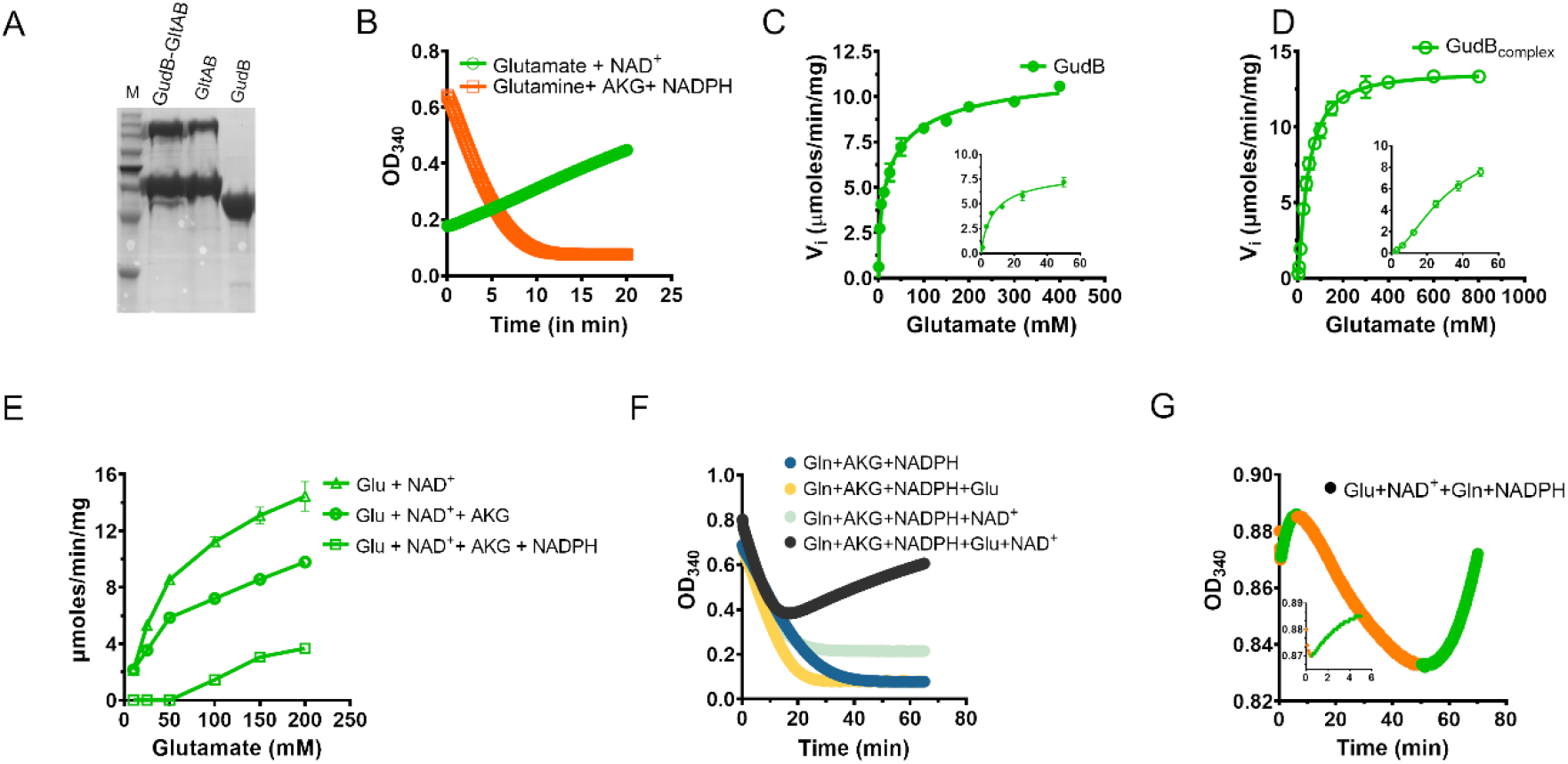
Enzymatic kinetics of the GltAB-GudB complex. **A)** SDS-PAGE analysis of the GltAB-GudB complex, and its individual components, GudB, and GltAB, used in these assays. The complex and stand-alone GltAB were purified by the pulldown of Strep-GltB from respective *B. subtilis* strains (see methods). GudB was purified by recombinant expression in E.coli. **B)** The GudB-GltAB complex exhibits either glutamate dehydrogenase activity (upon addition of glutamate and NAD+; green line, indicating a drop in absorbance at 340 nm due to NAD^+^ reduction) or synthetase activity (upon addition of AKG, glutamine and NADPH; orange line indicating an increase in absorbance due to NADPH oxidation). **C-D)** Initial velocity of the dehydrogenase reaction as a function of glutamate concentration. On its own, GudB displayed a K_M_ value of 7.3 mM for glutamate and negative cooperativity (C, Hill coefficient =0.57, Table 1). In complex with GltAB, the K_M_ rises to 138 mM with positive cooperativity (D, H=1.3). The inset in C and D shows the initial velocity at low concentrations of glutamate. Error bars indicate standard deviation of initial velocity measurement of two different batch of proteins and two technical replicates. **E)** Initial velocity of the dehydrogenase reaction in the GltAB-GudB complex, as is (Glu+NAD^+^), and in the presence of GltAB’s substrates (AKG, NADPH). Error bars indicate standard deviation of two independent measurements. **F)** Reaction progress curves of the GudB-GltAB complex with various substrate combinations: substrates of GltAB only (blue), substrates of GltAB plus glutamate (yellow), substrates of GltAB plus NAD^+^ (light blue), all substrates for both enzymes (black). **G)** Multiphasic progress curve displayed by the GudB-GltAB complex in the presence of substrates of both enzymes except AKG. The phases where GltAB predominates are shown in orange and those where GudB is dominant are shown in green. The inset shows the initial few minutes of the reaction where there is a rapid drop in absorbance followed by a gradual increase.

**Table 1:**
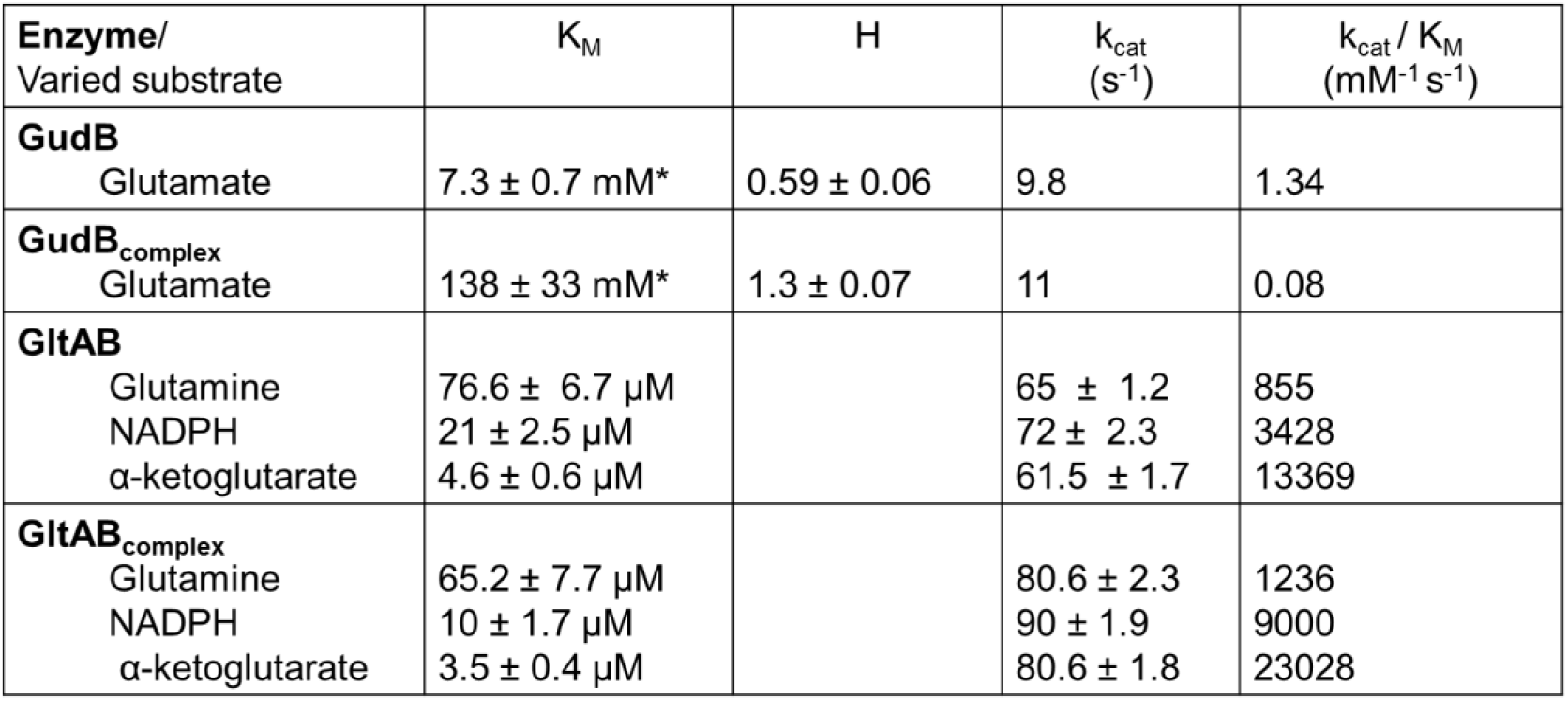
Steady state kinetic parameters of GudB, GltAB and the GudB-GltAB complex. Reactions were initiated by the addition of enzyme (5 µg of recombinant GudB, or 2.5 µg of the GudB-GltAB complex) and their progress was monitored by absorbance at 340 nm. The concentration of the fixed substrates were 4 mM of NAD^+^ for the dehydrogenase reaction, and 2 mM AKG, 5 mM glutamine, and 200 µM NADPH, for the glutamate synthase reaction. The reactions were carried out in 50 mM HEPES pH 7.9 at 25 °C. H is the Hills coefficient (measured for standalone GudB, and also for the GudB-GltAB complex, with glutamate as substrate). * The K_M_ values for GudB and GudB_complex_ were derived from fitting the data to an allosteric sigmoidal model. All data are presented as mean ± S.D of two independent measurements.

While the above measurements allowed us to derive kinetic parameters, the assay conditions do not reflect the cellular scenario where the substrates of both enzymes are present. Indeed, addition of GltAB’s substrates – AKG and NADPH, completely inactivates GudB at glutamate concentrations up to 50 mM (**Figure 3E**). The silencing effect seems synergistic as neither AKG nor NADPH exhibited a significant effect on their own. These results (**Figure 3C-E**) validate the hypothesis that GltAB’s binding silences GudB’s activity.

GltAB’s regulatory role was further manifested when its third substrate, glutamine, was added, unexpectedly resulting in multiphasic progress curves. The action of GltAB results in NADPH oxidation and is indicated by decrease in absorbance at 340 nm, while GudB’s activity that causes NAD^+^ reduction is manifested in an increase absorbance at 340 nm (**Figure 3B**). In principle, if these two enzymes that share substrates/products are present, a steady-state would be established. The progress curve should thus be monotonic, with the slope’s direction being either positive or negative depending on the relative catalytic efficiencies of the two enzymes and on substrate concentrations (Hart and Alon, 2013). For simplicity, consider the scenario where NAD^+^, NADPH, and glutamine, are well above saturation. Then, upon initiating the reaction if AKG is present at sufficient levels, the slope would be negative (GudB is suppressed and GltAB’s activity dominates). However, if glutamate is high, and AKG is sufficiently low, the slope would be positive – here, both enzymes might be active, but GudB’s rate would be higher resulting in net increase in 340 nm absorbance. Indeed, when AKG was present, despite an excess of glutamate, we observed the expected drop in absorbance indicative of GltAB activity (**Figure 3F**). Further, the slope of this first phase was identical to the one observed when GltAB activity was tested separately, suggesting that in this initial phase GudB is completely inactive due to the presence of AKG and NADPH (as shown in **Figure 3E**). However, this initial phase was followed by a second phase of increase in absorbance indicative of GudB activity. Indeed, when the reaction is initiated with all substrates except AKG, multiple oscillation cycles could be observed until most of the NADPH is consumed by GltAB resulting in GudB taking over (**Figures 3G** and **Figure S3 G-L**).

Oscillations need not involve transcription as with the circadian clock for example, and an oscillatory two-enzyme system has been described (Goldbeter, 1975; Masaki et al., 2013; Pálsson, 2009; Rossomando and Sussman, 1973). However, oscillations are not expected in a counter-enzyme pair as such. The rates of substrate binding, product release, and catalytic turnover, are typically in the sub-second range, and a steady-state would therefore be established with a constant ‘net’ rate (Hart and Alon, 2013). For oscillations to occur, a delay mechanism must take place (Blanchini et al., 2018; Hart and Alon, 2013; Radde, 2009; Wagner, 2005), namely, a slow, minute-time-scale, switch-over that will lead to “over-shooting” of one reaction before the next one takes over. Such a delay may be produced, for example, by a slow conformational change in GltA’s regulatory loop (*e.g.* via a proline cis-trans isomerization) and/or by cycles of dis-assembly and slow re-assembly of the complex. The latter would result in GudB over-producing AKG due to a delay in GltAB’s binding. That the oscillation frequency depends on the complex’s concentration, and in a non-linear way (**Figure S3 L-N**), is hinting at such a mechanism. The GudB-GltAB oscillatory kinetics may also have some intriguing physiological implications in biofilms, as elaborated in the Discussion section.

### Structure of the GudB-GltAB complex

Native mass spectrometry (native-MS) of the individual proteins provided preliminary information about the stoichiometry of the individual enzymes. As observed earlier (Gunka et al., 2010; Noda-Garcia et al., 2017), GudB is by itself a hexamer (**Figure 4A**). However, unlike its closest homologue with a known structure, Azospirillum GltAB, which is predominantly a hetero-dodecamer (Cottevieille et al., 2008; Swuec et al., 2019), *B. subtilis* GltAB is a hetero-dimer (**Figure 4A**). GltAB’s oligomeric state therefore resembles that of ferredoxin-dependent glutamate synthases found in photosynthetic organisms like plants or cyanobacteria (Kameya et al., 2007; Vanoni and Curti, 1999). The spectrum of the GudB-GltAB complex (**Figure 4B**) indicated a species with a mass of ∼1.6 MDa in the high m/z region, corresponding to six GudB subunits and six GltAB heterodimers (GudB_6_-GltA_6_B_6_) (**Figure S1B**).

**Figure 4.**
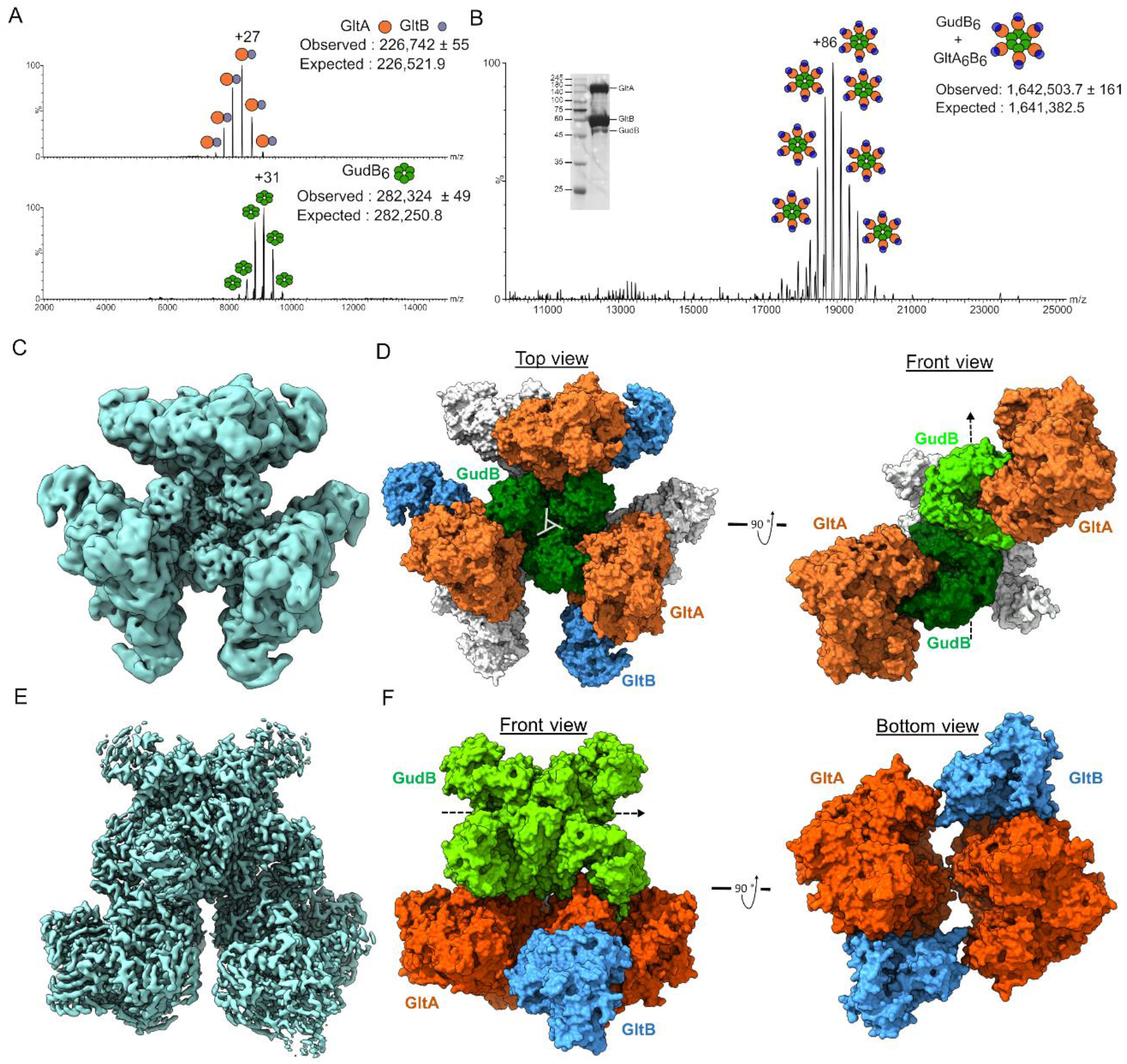
The stoichiometry, oligomeric state and atomic structure of the GltAB-GudB complex. **A)** Native-MS of standalone GltAB (top) and GudB (bottom). The oligomeric state as inferred from the mass is shown schematically besides each charge state: GltAB’s observed mass corresponds to a heterodimer (the difference is ascribed to truncations in GltB also observed in western blots, Figure S1B), while GudB’s mass indicated the expected hexamer. **B)** Native-MS of the GudB-GltAB complex. The observed mass corresponds to 6 copies of a GltAB heterodimer bound to the GudB hexamer (the difference is ascribed to truncations in GltB, as in panel a). The inset shows the SDS-PAGE of the protein sample used for native-MS and cryo-EM. **C)** Density map of the GudB_6_-GltAB_6_ complex generated after applying D3 symmetry. **D)** Model of GudB_6_-GltA_6_B_6_ complex. The structure on the left (top view) is shown with the dihedral (D_3_) rotation axis of GudB running orthogonal to the plane of the paper (the cross-section of the axis is shown as a triangle in the center of GudB, and the one on the right (side view), with the rotation axis running parallel to the plane of the paper (dashed arrow; GltB is omitted for clarity). **E)** The density map of GudB_6_-GltA_2_B_2_. **F)** On the left is shown the model of GudB_6_-GltA_2_B_2_ (front view) and right panel shows the same model from the bottom. From the bottom view, it can be seen that there are minimal interactions between the two GltA copies. GudB is omitted from the bottom view for clarity. Orange, GltA; blue, GltB; green and light green, GudB. Surface representation coloured white designate proteins behind the plane of view.

This complex is unusually large – its molecular weight is more than half of that of the ribosome, allowing us to apply CryoEM with the aim of validating the stoichiometry and the oligomeric state of GudB and GltAB in the complex, and to decipher how GltAB binding silences GudB. Using 12000 images, we obtained maps corresponding to the GudB-GltAB complex at a resolution of 3.9 Å (**Figure 4C, Figure S4 and Table S2**). Following refinement, we obtained a model of the complex that unambiguously revealed that the complex comprises a GudB hexamer and six GltAB heterodimers (overall correlation coefficient of 0.79; **Figure 4D**). Each GltAB heterodimer docks onto two subunits of GudB, with the interaction occurring solely through contacts with GltA. There are also no interactions between the GltAB hetero-dimers in the complex. Although the resolution of the map permitted assignment of the different chains, to reveal the mechanistic basis of GudB silencing by GltA, a map with higher resolution was obtained.

To increase the particle density, recombinant GudB was added to *B. subtilis* cell lysate before purifying the GudB-GltAB complex (**Figure S5A**). Preliminary cryo-EM single particle analysis resulted in four 3D classes representing varied number of GltAB hetero-dimers bound to the GudB hexamer (**Figure S5B**) in accordance with native-MS showing a range of GudB-GltAB complexes (**Figure S5A**). The four resulting maps all had a central barrel-like region which fits a hexameric GudB, with additional density that corresponds to GltAB heterodimers. Three minor classes, resemble the native complex with a GudB_6_-GltA_6_B_6_ stoichiometry with varying occupancy of the GltAB heterodimers. However, their preferred orientation biases prevented refinement to high resolution. The most abundant class corresponded to a GudB hexamer with two GltAB heterodimers (GudB_6_-GltA_2_B_2_). A high resolution map was prepared by collapsing these classes into a single class and refining with the preliminary GudB_6_-GltA_2_B_2_ map as the initial reference. This refined to high resolution (2.42 Å) with C2 symmetry (**Figure 4E**, **Figure S6 and Table S2**). The increased resolution permitted real space refinement using coot and ISOLDE, improving the model to map correlation to 0.83 (Figure 4F). Some weak density for the lower occupancy GltAB heterodimers remained around the periphery. This weak density suggests that although alignment focused on the best resolved GltAB chains, particles contributing to this map include additional partially assembled GltAB chains docked around the central GudB hexamer.

The higher resolution model confirmed the overall model observed in the native GudB_6_-GltA_6_B_6_ complex and allowed us to determine the precise contacts used to stabilize GudB in an inactive state (**Figure 4D**). It also comprises the highest resolution structure of GltAB. There appear to be no significant differences with the published 3.5 Å resolution structure of the hexameric GltAB from *Azospirillum brasiliense* (PDB ID: 6s6x) (Swuec et al., 2019). In Azospirillum GltAB, dimerization of GltA seems like a prerequisite for hexamer formation. Among other contacts, the GltA homo-dimer interface includes 13 putative hydrogen bonds. Examination of the corresponding residues in *B. subtilis* GltA suggested that 10 out of these 13 hydrogen bonds cannot be formed and in agreement with *B. subtilis* GltA not forming homo-dimers The higher resolution did permit unambiguous assignment of various co-factors that GltAB uses, viz., FAD, two 4Fe-4S clusters in GltB, and a 3Fe-4S clusters and FMN in GltA. These co-factors play a key role is shuttling electrons from the GltB-bound NADPH to 2-iminoglurate that is formed in the synthase domain of GltA (**Figure S7A**), thus allowing the reduction of the latter to give glutamate.

### The structural basis of GudB’s inhibition by GltAB

GudB interacts with GltAB extensively through the GltA subunit (**Figure 5A**) with few contacts between adjacent GltAB heterodimers (**Figure 4F, right panel**). GltA is a multi-domain protein that consists of an Ntn-amidotransferase domain, a central domain synthase domain, and the GXGXG domain (**Figure 5B**). The interactions with GudB occur primarily with GltA’s central domain that docks onto the active-site cleft of GudB. The latter resides between GudB’s cofactor (NAD^+^) and substrate (glutamate) binding domains (**Figure 5A**). The relative movement of these two domains results in “open” and “closed” conformations that are a key part of the catalytic cycle of glutamate dehydrogenases (Stillman et al., 1993). Cleft closure brings NAD^+^ close to glutamate thereby enabling the hydride transfer, as represented by the substrate bound structure of glutamate dehydrogenase from *Aspergillus niger (PDB: 5XVX)* (Prakash et al., 2018). Based on the distance between residues R280 and K122 as a proxy of the degree of closeness, GltA-bound GudB is in open state (d= 29.1 Å; **Figure 5C**). Another key observation is that each GltA subunit interacts with two adjacent GudB subunits (**Figure 5D**), also in agreement with GltA binding stabilizing GudB’s hexamer (**Figure S7B)**. Notably, the residues mediating this secondary interaction differ in RocG (underlined in **Figure 5E**) such that this interaction cannot be maintained and this in addition to few sequence differences between RocG and GudB (**Table S3**) could dictate the paralogue specificity of GltA.

**Figure 5.**
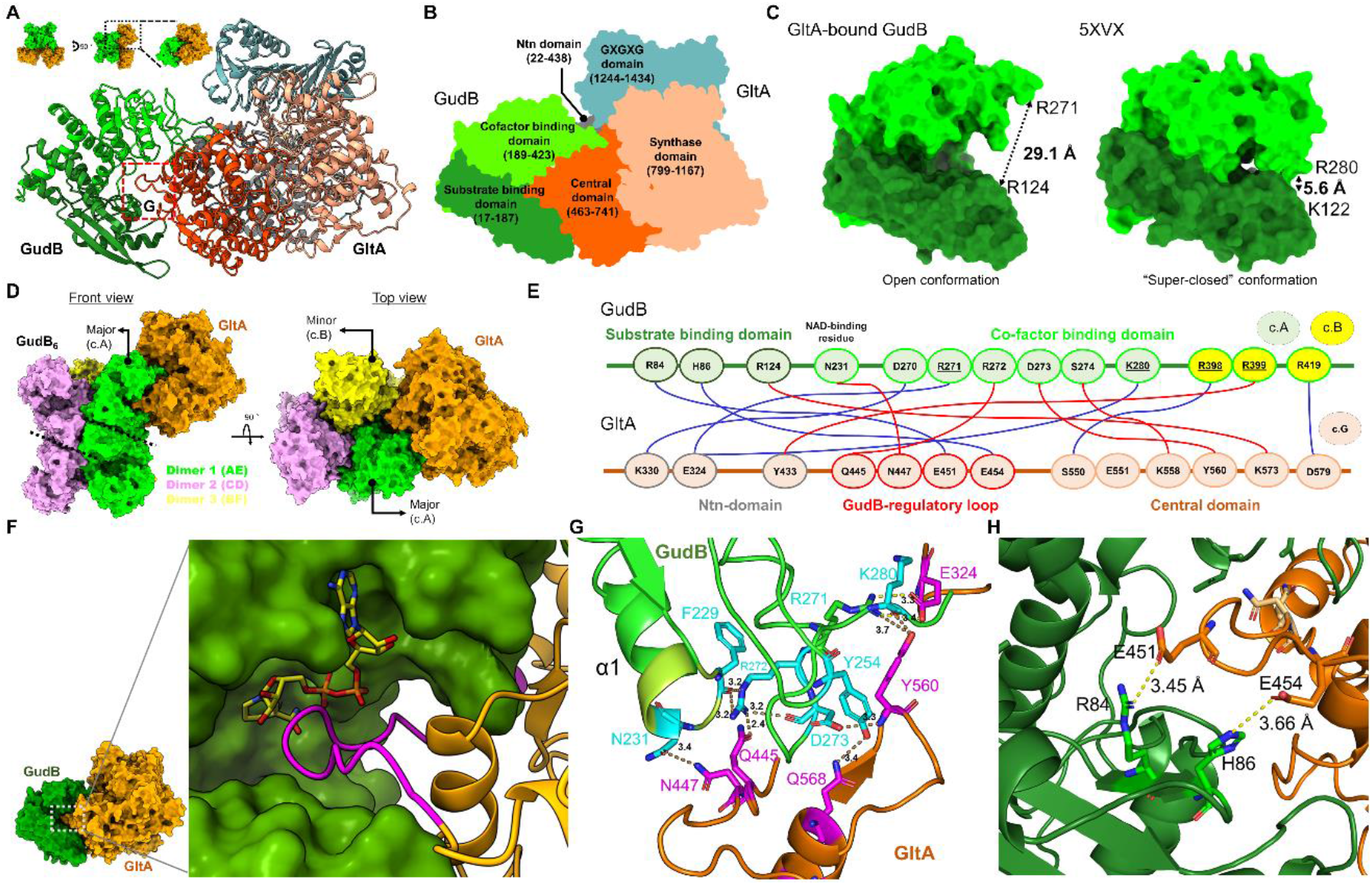
The structural basis for inhibition of GudB by GltA binding. **A**) Zoom-in model of the GudB-GltA interaction. The colors represent the different domains of GltA and GudB as detailed in the next panel. The inset shows the perspective of the model with respect to the model shown in **Figure 4F**. The dashed box (G) corresponds to the region containing residues from GudB and GltA forming a hydrogen bonding network. This is shown in detail in panel G. **B**) GltA consists of Ntn amidotransferase (Ntn) domain (grey), a central domain (orange), a synthase domain (sand) and a GXGXG domain (metal blue). The central domain of GltA (orange) interacts with residues from both the substrate (green) and co-factor binding domain of GudB (light green). **C**) GltAB binding captures GudB in an “open” state with the distance of 29.1 Å between residues R280 (in the co-factor binding domain, shown in light green) and K122 (in the substrate binding domain, shown in green). These residues are equivalent to R271 and R124 in the model of a “super-closed” glutamate dehydrogenase (PDB 5XVX) (Prakash et al., 2018) (right panel). **D**) GudB’s hexamer comprises a trimer of dimer; GltA (in orange) interacts with two protomers of GudB (major and minor) from two different dimers (green and yellow, respectively). The front view shows the major interacting GudB protamer, and the top view the minor one. The coloring pattern of the three dimers and the corresponding chain ID’s are also shown. The dimer interface in two of the dimers (1 and 2) is shown as dashed lines. **E**) Key interactions between GltA (light orange coloured ovals) and GudB (light green and yellow colored ovals; salt bridges in red and hydrogen bonds in blue). For GudB, the outline color of each oval corresponds to the domain (substrate/cofactor-binding) to which the interacting residue belongs to. Residues from chain A are shown in green oval and chain B are shown in yellow. F) Zoom-in on GudB (surface display, in green) and the interacting loop of GltA (magenta; the remaining structure of GltA is in orange). NAD^+^ (in yellow sticks) is modelled into the active site cleft of GudB based on superposition of GltA-bound GudB to 1V9L (open state). The steric overlap of GltA’s loop with NAD^+^ would be even higher in the closed state of the enzyme. **G**) Zoom-in on the boxed region shown in panel A, depicting the hydrogen bonding network between residues of GltA (magenta) and the co-factor binding domain of GudB (cyan). Notably, GudB’s N231, which is located at the tip of the phosphate binding loop (and binds the NAD^+^’s phosphate groups) is bound to GltA’s N447, and thus directly interferes with NAD^+^ binding. **H**) The interaction of GltA’s E451 and E454 with residues of the substrate binding domain of GudB (R84 and H86).

The primary interacting subunit (“major”) has its active site blocked as described above. The neighboring subunit interacts with a much smaller surface area outside the active site (**Figure 5D, top view**). Despite a large interface area (1931 Å^2^ of GudB and 1817 Å^2^ of GltA), there are relatively few specific interactions (6 salt bridges and 6 hydrogen bonds) as shown in **Figure 5E** (a complete list of interactions is provided in **Table S3**). The key interaction between GudB and GltA involves a long loop, “the GudB regulatory loop” that connects the Ntn- and central-domains of GltA (residues 445-455). This loop extends across the active site cleft of GudB. Residues from the loop interact with both the cofactor- and substrate-binding domains of GudB (**Figure 5F-G**). Foremost, a network of hydrogen bonds is observed between GltA residues and the cofactor binding domain of GudB (**Figure 5F**), which in turn stabilizes multiple loops in GudB. Notably, clear density for these loops is seen in the complex that was not seen in the apo-GudB crystal structure (Gunka et al., 2010) or in the GudB subunits in the complex that are not bound to GltA. The ordering of the cofactor binding domain of GudB by binding to GltA is also reflected in the lower average b-factor of GltA-bound GudB compared to the unbound one (**Figure S7C**). The most notable contact is between the tip of α1 and residue N231 of GudB’s NAD^+^ binding domain. In dehydrogenases of the Rossmann fold, the tip of α1 is typically used to bind the di-phosphate group of NAD^+^. Consequently, GltA binding precludes NAD^+^ binding to GudB. Moreover, as shown in **Figure 5H**, GltA’s regulatory loop occupies a specific position in the active site cleft of GudB which is otherwise occupied by the co-factor in the closed (and catalytically active) conformation of the enzyme. Thus, stabilization of the open conformer, and blocking NAD^+^ binding by steric hindrance, seem to be the key to GudB’s inhibition.

### GudB-GltAB interaction in biofilm formation

In planktonic growth, the role of GudB-GltAB complex formation is silencing of GudB under glutamate poor growth condition (**Figure 2A** and **2E**). What could be its physiological significance for biofilm growth? Bacillus is present as biofilms in its natural environment, and glutamate metabolism plays a key role in biofilm development (Hassanov et al., 2018; Pisithkul et al., 2019; Zhang et al., 2015). Accordingly, in the laboratory, *B. subtilis* biofilms are usually grown in glutamate rich media containing glycerol as the carbon source (MSGG). However, unlike planktonic growth, biofilm growth on solid agar presents spatiotemporal contraints – cells in the interior depend on the peripheral cells for glutamine, while the peripheral cells depend on the interior cells for ammonia (Liu et al., 2015). The GudB-GltAB complex could be central to this division of labour.

We initially examined the morphology of biofilms that are devoid of either GudB, GltA, or GltB. Unlike for planktonic growth (**Figure 2A-D**), for wild-type like biofilm growth and morphology, all three proteins seem to be important (**Figure 6A**). *ΔgudB* produced rapidly growing biofilms (**Figure S8**) with large “channels” running from the periphery to the interior. The Δ*gltA* and Δ*gltB* strains also showed biofilm morphologies that differ from wild-type – while Δ*gltB* strain had wrinkles restricted to interior, Δ*gltA* biofilm did not have any wrinkles. Since the enzymatic function, glutamate synthesis, is lost in the absence of either GltA or GltB, the different phenotypes are in agreement with the additional role for GltA in regulating GudB.

**Figure 6.**
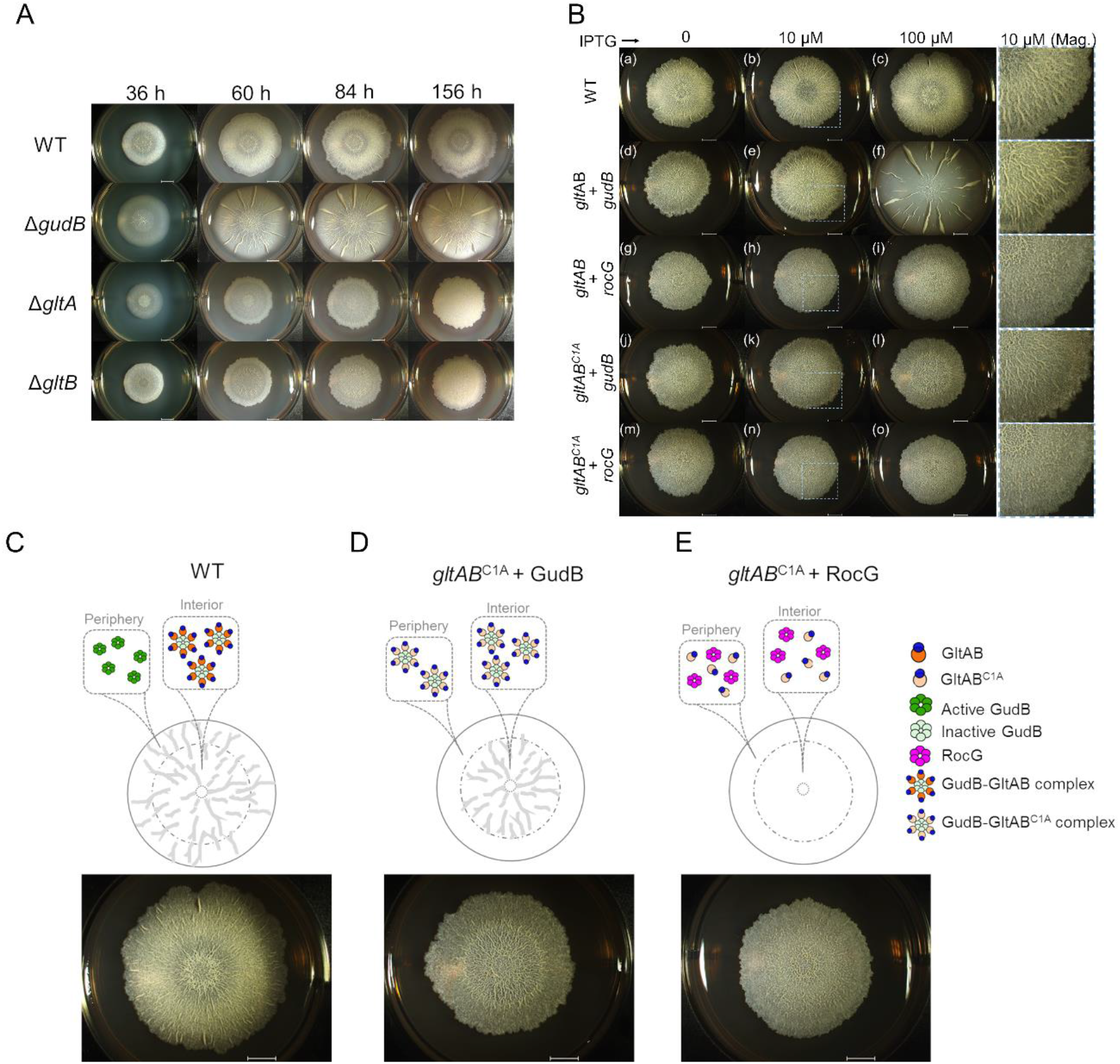
GudB-GltAB interaction is important for biofilm formation. **A)** Representative images of biofilms of wild-type and mutant *B. subtilis* strains on glutamate-glycerol (MSGG) agar medium, monitored over a period of ∼7 days. As indicated by their knockouts, both Glt subunits (GltA and B), and GudB, are essential for wild-type like biofilm morphology **B)** Comparision of biofilm formation by wild-type and mutants expressing GltA under an IPTG-induced promoter, at varying IPTG concentrations (0-100 µM). Wild-type (a-c); IPTG-induced GltA plus GudB (d-f), or RocG (g-i); IPTG-induced inactive GltA mutant, GltAB^C1A^ plus GudB (j-l), or with RocG (m-o). Wild-type GltAB and the GltAB^C1A^ were expressed from an IPTG-inducible hyperspank promoter, while GudB and RocG were under *gudB’s* promoter. Wild-type’s biofilms are not affected by the presence of IPTG (a-c). The strain expressing GltAB from an IPTG-induced promoter forms altered biofilms with no IPTG (d), wild-type-like biofilms with low concentration of IPTG (e), while overexpression resulted in biofilms similar to those of the *ΔgudB* strain (f). Co-expression of RocG and GltAB resulted in biofilms with no wrinkles (g-i). Expression of the inactive GltAB^C1A^ with GudB restores the morphology of the biofilm’s interior, in an IPTG-dependent manner (j-l); in contrast, co-expression of GltAB^C1A^ with RocG does not (m-o). The scale bars in all images correspond to 4 mm. Magnified sections are shown in the last column of panel B. **C-E)** A summary of the biofilm disruption phenotypes and a proposed model that accounts for them (see text). The biofilm images correspond to c, l and o (from left to right in that order) in panel B.

Next we examined the role of GudB-GltAB complex formation using the strains that express either wild-type GltAB, or its inactive mutant (GltAB^C1A^), from an inducible promoter (see GltAB negatively regulates GudB activity in vivo above). In the strain having GltAB under an inducible promoter, induction at low levels (10 μM IPTG) was sufficient to restore biofilm formation to wild-type levels (**Figure 6B** (e); wild-type biofilms were not affected by addition of IPTG; **Figure 6B** (a-c)). However, overexpression of GltAB (100 µM IPTG), resulted in Δ*gudB* like biofilm morphology (**Figure 6B(f), and Figure S8B** for enlarged images) consistent with silencing of GudB by GltAB. To tease out the differences between the catalytic and regulatory role of GltA, we examined the biofilms of cells expressing the inactive GltAB^C1A^. Here, wild-type-like growth was restored, but only in the interior of the biofilm (**Figure 6B** (j-l) and **Figure 6C**), suggesting that silencing of GudB by GltAB is crucial for the interior of the biofilm. In contrast, if the GudB silencing occurs in the peripheral cells, the biofilm exhibits no wrinkles in the periphery (compared to wild-type; **Figure 6C**). This result is also in agreement with GltAB’s expression being the highest in the interior of the wild-type biofilm (Liu et al., 2017). Hence, it appears that regulation of GudB’s activity by GltAB binding is crucial for wild-type like biofilm formation. Indeed, when the above-described wild-type-and inactive-*gltAB* genes were expressed at the background of RocG, the effects differed completely from those seen at the GudB background (**Figure 6B** (g-I, m-o)). Foremost, in the presence of RocG, expression of GltAB^C1A^ had no effect and the biofilms completely lacked wrinkles (**Figure 6C**). This result is consistent with RocG having the same level of enzymatic activity as GudB, yet not being suppressed by GltAB binding (see ‘the interaction with GltAB is paralog specific’ above). Overall, these results show that while enzymatically functional GudB and GltAB are important for biofilm growth, so is the silencing of GudB by the formation of GudB-GltAB counter-enzyme complex.

## Discussion

Our search for the regulatory mechanism of GudB, a core catabolic glutamate dehydrogenase which is constitutively expressed, yielded an unexpected answer. GudB, which is involved in glutamate catabolism, forms a transient complex with its counter-enzyme GltAB, that catalyses glutamate synthesis. The main purpose of this association seems to be regulatory.

Most biochemical transformations are in principle reversible. However, *in vivo*, many of these are effectively unidirectional due to thermodynamic and kinetic constraints. In addition, many enzymes evolved towards unidirectional catalysis, including oxidoreductases that utilize NAD^+^ to drive the reaction in the oxidative direction (*e.g.* GudB) or NADPH to direct reduction (*e.g.* GltAB).

Consequently, cells contain multiple counter-enzymes that catalyze opposite transformations, or (note that the reactions themselves usually differ, as is the case with GltAB and GudB, but the primary substrate/product are the same as shown in **Figure 1A**). It yields to reason that counter-enzymes would be individually regulated, as their simultaneous operation results in futile cycles (zero net outcome and waste of energy). However, regulation via counter-enzyme pairs offers unique properties that cannot be obtained by independent regulation of the individual enzymes, such as ultrasensitivity, robustness, and temporal signaling pulses (Hart and Alon, 2013). The opposing enzymes can be separate proteins, or fused on the same polypeptide chain (bifunctional enzymes) (Hart and Alon, 2013; Straube, 2013). Such pairs involved in signaling have been characterized. These usually mediate cycles of introducing and erasing a covalent post-translational modification (*e.g.* phosphorylation-dephosphorylation, acetylation-deacetylation etc.). However, in core metabolism, although multiple counter-enzyme pairs exist, regulation through counter-enzyme pairs has not been described to our knowledge. Given the GudB-GltAB example unraveled here, we speculate that other counter-enzymes pairs might exist in core metabolism.

Our findings also highlight a new feature of complexes in regulating the activity of core metabolic enzymes. The traditional view of the cell, or its cytoplasm, as a homogenous environment in which enzymes and metabolites diffuse freely is being revised. The role of enzyme clustering, of transient, ad-hoc associations of enzymes that share substrates or intermediates, so-called metabolons, is increasingly apparent (Pareek et al., 2020; Zhang and Fernie, 2020). Metabolons have been primarily associated with compartmentalization of reactions, channeling of substrates and intermediates between enzymes, and prevention of their leakage. The GudB-GltAB association is an example of the key role that enzyme complexes can play in regulating enzyme activity, and specifically in regulating counter-enzymes.

Under planktonic growth, regulation regards the silencing of GudB by association to GltAB. Silencing is critical in conditions where glutamate is being made rather than broken, typically with carbon sources such as glucose that are converted to AKG (by the action of glycolysis and a set of TCA enzymes). However, the GudB-GltAB complex seems to offer a more elaborate regulatory mode that may be of particular importance in biofilms. During biofilm growth, glutamate is readily available to the peripheral cells while the interior cells experience glutamate starvation (Liu et al., 2017). This spatial constraint demands that the catabolic GudB is active in the peripheral cells, while the anabolic GltAB is active in the interior. While GltC-dependent transcriptional control of *gltAB* (Picossi et al., 2007) ensures that GltAB is expressed in cells starved for glutamate, the GudB-GltAB counter-enzyme complex formation ensures that GudB (that is constitutively expressed) is active in the periphery and silenced in the glutamate-starved interior cells (**Figure 6C**). Accordingly, constitutive expression of GltAB^C1A^ and RocG that cannot form the complex completely disrupts the biofilm morphology (**Figure 6E**). In contrast, constitutive expression of GltAB^C1A^ along with GudB, silences GudB via complex formation in both interior and peripheral cells resulting in biofilms that look like wild-type in the interior but not in the periphery (**Figure 6D**). This unique mode of glutamate dehydrogenase regulation may also relate to *B. subtilis* biofilms exhibiting oscillations in their growth rates (Liu et al., 2015). These oscillations have been attributed to metabolic codependence between cells in the interior and in the periphery of the biofilm. Modelling these oscillations revealed that they depend on a highly cooperative glutamate dehydrogenase (Hill coefficient of 12), in effect, a hair-triggered enzyme (Bocci et al., 2018). While modelling did not address a specific enzyme, GudB is the key glutamate dehydrogenase in wild-type *B. subtilis*, and its paralog, RocG, was shown to have no effect on oscillations (Liu et al., 2015). The oscillatory kinetics exhibited by the GudB-GltAB complex (**Figure 3G**) likely relate to the biofilm oscillations, and future work may establish a direct link between these two phenomena.

Oscillations in signaling reactions have been noted, in relation to the circadian clock, but also in cAMP signaling (Goldbeter, 1975). However, to our knowledge oscillations in the activity of metabolic enzymes have not been seen. In cAMP signaling, oscillations are induced by the action of a multi-enzyme system (adenylate kinase, ATP pyrophosphate hydrolase, intra- and extracellular phosphodiesterase) along with cAMP receptor, that together mediate cAMP-dependent cell-cell aggregation in *Dictyostelium discoideum* (Gregor et al., 2010; Pálsson, 2009; Rossomando and Sussman, 1973). These oscillations are driven by the products of both enzymes (cAMP, and AMP, respectively) activating the other enzyme. In the GudB-GltAB complex oscillations seem to be driven by the inhibition of one enzyme (GudB) by the other (GltAB) and also by physical association between the two enzymes (which does seem to be part of the cAMP oscillations mechanism). Future work may therefore enable a deeper understanding of the GltAB-GudB complex, and also of the potential correlation between GudB’s regulation and biofilm oscillations.

## Acknowledgments

We thank Kesava Phaneendra Cherukuri for help in synthesis of DSG, Alexander Leytens for assistance in cloning, and Yuval Kushmaro for screening different crosslinkers. We are grateful to Meital Kupervaser and Yishai Levin from The De Botton Protein Profiling institute of the Nancy and Stephen Grand Israel National Center for Personalized Medicine, Weizmann Institute of Science, for the proteomics analysis. We thank Prof. Uwe Sauer for his critical feedback on an earlier version of this manuscript. We thank Dr. Felix Jonas for valuable comments on the manuscript and Dr. Brian Ross for proofreading the manuscript. Funding by the Israel Science Foundation Grant No. 2575/20 is gratefully acknowledged. M.S. is the incumbent of the Aharon and Ephraim Katzir Memorial Professorial Chair. The research of S.V is supported by the Clore Israel Foundation. J.S.F is supported by NIH GM123159 and D.J.L is supported by NIH F32 AI148120. D.S.T is the incumbent of the Nella and Leon Benoziyo Professorial Chair.

## Author Contributions

D.T. conceptualized the study and supervised all research; V.J. performed all experimental work unless otherwise stated; S.V. performed the native mass spectrometry and M.S. supervised this analysis; N.E., acquired the Cryo-EM data, and D.J.L., V.J. and N.E. performed the Cryo-EM data processing and model building and was supervised by J.S.F. V.J. and D.T wrote the manuscript and all authors reviewed the data and the manuscript.

## Methods

### DNA manipulation

All constructs used for genome modification were generated using Gibson assembly. The starting fragments were generated using Q5^®^ High-Fidelity DNA Polymerase (New England Biolabs) and NEBuilder^®^ HiFi DNA Assembly Master Mix (New England Biolabs) was used for assembly. The linear DNA fragments for *B. subtilis* transformation were generated by digestion of the plasmids (using a single restriction enzyme) or by PCR using the Gibson assembly reaction as template. The fragments had recombination homology arms ranging anywhere from 1.5 to 4 kb. All the primers used for the study are listed in **Appendix Table 2**. Genetic modification of *B. subtilis* strains were performed based on natural competence: The parental strains were streaked on an LB-agar plate and grown overnight at 37 °C. A singly colony was then used to inoculate 1 ml of modified MC medium (Wilson and Bott, 1968) containing 0.5% glucose, 1.4% K_2_HPO_4_, 0.6 % KH_2_PO_4_, 30 nM sodium citrate tribasic dehydrate, 0.2 % casein hydrolysate, 84 mM ammonium iron (III) citrate, and 3 µM MgSO_4_. Addition of tryptophan (0.005%) and histidine (0.005%), improved the transformation efficiency. The culture was incubated at 37 °C in a roller drum shaker for 3.5 hrs. Linear DNA/ genomic DNA (100 ng -1 µg) was added to 300 µl of this culture and growth was continued for another 3 hrs. The positive transformants were selected either on an antibiotic, or in nutrient selection media (for wild-type GltAB constructs). All transformants were verified by diagnostic PCR’s followed by DNA sequencing.

### Media and growth conditions

*B. subtilis* strains were grown in LB, or in MS medium (5 mM potassium phosphate, 100 mM MOPS pH 7.1, 2 mM MgCl_2_, 700 µM CaCl_2_, 50 µM MnCl_2_, 50 µM FeCl_3_, 1 µM ZnCl_2_, 2 µM thiamine; adapted from (Branda et al., 2001). The medium was supplemented with appropriate carbon/nitrogen source (glutamate, histidine, arginine, proline) or with carbon source (glucose, glycerol) plus ammonium sulfate, all at a final concentration of 5 g/L. Media were supplemented with antibiotics when required: kanamycin (10 µg/ml), tetracycline (5 µg/ml), spectinomycin (100 µg/ml) or a mix of lincomycin (25 µg/ml) plus erythromycin (2 µg/ml). Growth profiling of *B. subtilis* strains in liquid MS medium was performed in a 96 well plates and absorbance at OD_600_ was measured using a multi-well plate reader (Eon, Biotek). Typically, 3 µl of culture with an OD_600_ of 0.8-1.2 was used as a starter to inoculate a well containing 200 µl fresh medium. Replicates were performed parallely in the same plate.

For biofilm growth, the MS media (supplemented with appropriate C/N source) was solidified using 1.5 % agar (w/v). The plates were then dried for 6-8 hrs in a laminar hood. For biofilm growth, cells stocks were streaked on to an LB agar plate and incubated overnight at 37 °C. From the confluent part of the plate, cells were swiped using a sterile loop to inoculate a 5 ml LB medium, then grown at 37 °C shaker for 2-3 hrs (OD_.600_ of 0.8-1.2). Following this the cells were washed twice with 1 X PBS. The cultures were normalized based on OD_600_ and a 3 µl aliquot of a 0.6-1.0 OD_600_ culture was placed at the center of the plate for biofilm growth, and the plates were incubated at 30 °C. Photographs of the biofilms were acquired using Stereo Discovery V20 microscope with Objective Plan Apo S 0.5× FWD 134 mm or Apo S 1.0× FWD 60 mm (Zeiss) attached to an Axiocam camera. Data were captured and analyzed using ZENpro AxioVision suite software (Zeiss, Oberkochenm, Germany).

### Immunoprecipitation and mass spectrometry

Anti-GudB antibodies were generated in rabbits using recombinant GudB purified from *E. coli* as antigen. The IgG fraction from these polyclonal sera was purified using a Protein A column. The purified antibodies were covalently linked to CNBr beads (Kavran and Leahy, 2014a) and this resin was used for the pulldown of GudB from *B. subtilis* cell lysates. Briefly, wild-type *B. subtilis* (NCIB 3610) was grown in 500 ml MS medium supplemented with either glucose-ammonia or histidine. Cells were harvested by centrifugation (6100 rcf) and resuspended in 30 ml of lysis buffer (100 mM Tris. HCl, 150 mM NaCl, EDTA-free protease cocktail inhibitor (1:200, Abcam), 2 mM DTT). The cells were lysed by passing cell suspensions 3-4 times through a french press at a pressure of 15 psi. The debri was removed by centrifugation (11,648 rcf) and the resulting supernatant was mixed with the anti-GudB antibodies resin and incubated for 1 hr. After rinsing the beads, the bound proteins were eluted using a low-pH glycine buffer (100 mM, pH 2.8). Eluates were neutralized by collecting them directly into 2 M Tris buffer pH 8.5, pooled, and lyophilized. The eluted proteins were resolved on SDS-PAGE and stained with silver nitrate using established protocols (Kavran and Leahy, 2014b).

For MS-based identification of proteins, the dried eluates were resuspended in 5% SDS in 50 mM Tris pH 7.4 and subjected to tryptic digestion using an S-trap (Elinger et al., 2019). The resulting peptides were analyzed using nanoflow liquid chromatography (Acquity M-class) coupled to high resolution, high mass accuracy mass spectrometry (Q Exactive HF). Each sample was analyzed on the instrument separately, in a random order, discovery mode. Raw data were processed with MaxQuant v1.6.0.16, and searched with the Andromeda search engine against the *B. subtilis* proteome database appended with common lab protein contaminants, with the following modifications: Carbamidomethylation of cysteine as a fixed modification and oxidation of methionine as a variable one. The LFQ (Label-Free Quantification) intensities were calculated and used for further calculations using Perseus v1.6.0.7. Decoy hits were filtered out, as well as proteins that were identified on the basis of a modified peptide only.

### Protein expression and purification

Recombinant GudB and RocG were purified as described (Noda-Garcia et al., 2017). Briefly, the plasmids pET28_Strep_GudB and pET28_Strep_RocG encoding *B. subtilis* GudB and RocG respectively, were used for expression in the *E. coli* strain BL21 star/DE3 (pGRO7). Typically, the cells were grown in 1 L of terrific broth supplemented with kanamycin (50 µg/ml) and chloramphenicol (34 µg/ml) to an OD_600_ of 0.6-0.8 and protein expression was induced by the addition of 100 µM IPTG. Post-induction, the cultures were grown overnight at 20 °C with shaking (200 rpm). Cells were harvested by centrifugation (6100 rcf), resuspended in lysis buffer (100 mM Tris. HCl, 150 mM sodium chloride, protease cocktail inhibitor) and lysed using french press at 15 psi. The clarified cell lysate was applied on to a Strep-Tactin column, unbound proteins were removed by washing with 5 CV of lysis buffer, and the target protein was eluted using 10 mM desthiobiotin in lysis buffer. The eluted protein was then dialyzed overnight in storage buffer (50 mM HEPES pH 7.9); aliquots were flash frozen in liquid nitrogen and stored at -80 °C freezer for later use.

The GudB-GltAB complex was purified directly from *B. subtilis* strain in which GltB is expressed as a Twin-strep tagged protein (GltB-TS). For the purification of GltAB alone without GudB, the Δ*gudB* version of the strain expressing GltB-TS was used. A higher yield of the complex was obtained upon using strains containing *gltAB* genes under IPTG inducible-hyper spank promoter. GltAB contains Fe-S clusters that are essential for its activity; to prevent cluster oxidation, all buffers used for the purification were degassed first by applying vacuum and then purging the solution with nitrogen for ∼40 min. additionally, the buffer was supplemented with 5 mM DTT to maintain reducing conditions. For purification, the relevant *B. subtilis* strain was grown in 1-2 L MS medium containing glucose-ammonia to an OD_600_ of 1.5-2. The cells were harvested by centrifugation, resuspended in lysis buffer (100 mM Tris. HCl, 150 mM sodium chloride, protease cocktail inhibitor, 2 mM DTT) and lysed using a French press. The clarified cell lysate was either directly applied on to a Strep-Tactin column or in case of GudB enriched sample, the lysate was incubated with lysate from the *E. coli* overexpressing 6X His-tagged GudB. The column was washed with 5 CV of lysis buffer and the bound proteins were eluted using lysis buffer containing 10 mM of desthiobiotin. Elution aliquots were immediately flash frozen and stored in -80 °C. Quantification of the eluates were performed using Bradford assay (Bio-Rad). The concentration of individual proteins in the complex was determined by densitometric analysis on ImageJ2 (Rueden et al., 2017).

Paralogue specific binding of GltAB was tested by pulldown of GltB-TS from a strain expressing either RocG or GudB. Cell lysates from the respective strains were applied onto a Strep-Tactin column, unbound proteins washed, and the bound proteins eluted using 10 mM desthiobiotin in lysis buffer. The proteins in the eluate were separated on an SDS-PAGE and Coomassie stained. For the reciprocal pulldown, purified recombinant 6XHis RocG, or GudB, were added to *B. subtilis* cell lysates containing GltAB-TS. Bound proteins were was pulled down as above. Interactions were also tested *in vitro*, by adding either recombinant 6X His tagged GudB/RocG to *B. subtilis* cell lysates containing GltAB-TS. The lysate was applied on to a Ni-NTA column and the column was washed with 10 CV of wash buffer. Proteins bound to the column were eluted using 500 mM imidazole in lysis buffer. The eluates were resolved on SDS-PAGE and the gel was visualized using Coomassie staining.

### Crosslinking and western blotting

Disuccinimidyl glutarate was synthesized using established protocols. Briefly, glutaric acid (2.0 g, 1.0 eq) and N-hydroxy succinimide (3.8 g, 2.2 eq) were added to an oven dried round bottom flask containing 30 ml of dimethylformamide (DMF) and kept on ice. Then, EDC.HCl (6.4 g, 2.2 eq) was added to the flask one ice, and the reaction was subsequently stirred at room temperature overnight. The reaction mixture was then poured onto an ice-cold solution of 1 N HCl (∼10 volumes) and stirred for few minutes. The resultant solution was filtered and the precipitate was rinsed twice with ice-cold isopropanol. The obtained solids were dried under vacuum and further lyophilized to remove traces of water. The lyophilized powder was stored at – 20 °C. For crosslinking, wild-type *B. subtilis* strain NCIB 3610 was grown in 10 ml of MS-medium supplemented with appropriate C/N source to an OD.600 of 1-1.5. Cells were harvested by centrifugation and washed with 1 X PBS. The cells were resuspended in 500 µl of 1 X PBS. DSG was freshly dissolving in anhydrous DMSO to 250 mM, and added to the cell suspension to a final concentration of 0.5 – 2 mM. The cells were incubated in a tumbler, at room temperature, for 30 min unless otherwise specified. The crosslinking reaction was stopped by the addition of 2 M Tris pH 8.5 (to a final concentration of 150 mM) and incubated for another 15 min. The cells were harvested by centrifugation and the cell pellets stored at -20 °C.

For western blotting, the pellets of DSG treated/untreated cells were resuspended in 1 X SDS-PAGE loading dye and heated at 95 °C for 15 min. The cell debris was removed by centrifugation and the supernatant was analyzed on 8 or 12 % SDS-PAGE gel. The loading was normalized based on OD_600_ of the culture before crosslinking. The resolved proteins were transferred onto a PVDF membrane and blocked with 5 % skim milk solution in 1 X PBS at room temperature for 1 hr. Anti-GudB antibody (Protein A purified) was added (1: 50,000) to the blocking solution and the membrane was incubated for 1 hr. Following this, the membrane was subjected to 3 wash cycles, 5 min each, in washing buffer containing 0.01 % Tween (in 1 X PBS). Following the washes, HRP-conjugated anti-rabbit IgG, was added (1: 50000) and incubated with shaking for 1 hr. The blot was washed again and the protein bands were visualized using the ECL system and images acquired in Amersham imager 680, General electric company.

### Enzyme kinetics

Glutamate dehydrogenase and glutamate synthase activities were monitored by a change in absorbance at 340 nm. While the dehydrogenase reaction involves NAD^+^ reduction and hence increase in absorbance as the reaction proceeds, the synthetase reaction involves NADPH oxidation and decrease in absorbance. All kinetic assays were performed in multiwell plates suitable for measuring in UV range (Microplate UV/VIS 96F, Eppendorf). Reactions were performed in 50 mM HEPES buffer pH 7.9, at total volume of 200 µl, and were initiated by enzyme addition (5 µg of recombinant GudB and 2.5 µg of GudB-GltAB complex). Steady-state kinetic parameters for the enzymes were derived from initial velocity measurements where one substrate was titrated keeping all other substrates saturating. For GudB, the saturating concentration for glutamate was 200 mM, and for NAD^+^, 4 mM. For GltAB, the saturating concentrations of glutamine was 100 mM, AKG, 2 mM, and NADPH 200 µM. In addition to these substrates, all reactions involving GltAB included 5 mM MgSO_4_ and 5 mM DTT. The data were fitted to Michaelis-Menten or to an allosteric sigmoidal model in GraphPad Prism (9.1.0).

### Native mass spectrometry

All native-MS measurements were preformed using Q-Exactive UHMR instrument (Thermo Fisher Scientific, Bremen, Germany). Prior to the analysis, samples were buffer exchanged into 150 mM of ammonium acetate pH 8 using Bio-Spin P6 columns (BioRad) and the final protein concentrations were adjust to 1-5 µM. The following parameters were used for MS1 experiment: capillary voltage 1.1 k, desolation voltage -100 V, source fragmentation 0 V, HCD energy 50 V, trapping pressure 7, corresponding to HV pressure of 2.04 × 10^-4^ mbar. All spectra were recorded at resolution of 6250 at 800 m/z and analyzed using Xcalibur and, Masslynx4.2 (Waters) data analysis software. Spectra are shown without any smoothing, and the instrument were externally mass-calibrated using a 2 mg/mL cesium iodide solution.

### Cryo-EM sample preparation

Both GudB_6_-GltA_2_B_2_ and GudB_6_-GltA_6_B_6_ protein samples were prepared at a protein concentration of 0.5 mg/ml and applied to glow discharged R 2/2, 300 mesh, grids with an additional thin layer (∼2 nm) of continues carbon (Quantifoil). The GudB_6_-GltA_2_B_2_ complex solution was applied to the grid in a single 2.5 µl drop, followed by 30 sec incubation before blotting. The GudB_6_-GltA_6_B_6_ sample was applied in two consecutive drops of 6 and 3 µl in order to increase complex concentration on the grid. The first drop was blotted manually, followed by application of the second drop and automated blotting in the plunger. Both samples were plunge frozen in liquid ethane cooled by liquid nitrogen using a Vitrobot plunger (Thermo Fisher Scientific). The plunger was set to 3 sec blotting time and 100% humidity.

### Cryo-EM data collection

Data collection and statistics are summarized in **Table S2 and Figures S4** and **S6**. Cryo-EM data were collected on a Titan Krios G3i transmission electron microscope (Thermo Fisher Scientific) operated at 300 kV. Movies were recorded on a K3 direct detector (Gatan) installed behind a BioQuantum energy filter (Gatan), using a slit of 15 eV. Movies were recorded in counting mode at a nominal magnification of 105,000x, corresponding to a physical pixel size of 0.82 Å. The GudB_6_-GltA_6_B_6_ data set was collected at a dose rate of 21.4 e^-^/pixel/sec and total exposure time of 1.5 sec, resulting in an accumulated dose of 47.7 e^-^/Å^2^. The GudB_6_-GltA_2_B_2_ data set was collected at a dose rate of 20.5 e^-^/pixel/sec and total exposure time of 2 sec, resulting in an accumulated dose of 61 e^-^/Å^2^. Each movie was split into 60 frames, and the nominal defocus range was -0.8 to -2 μm. Each movie was split into 45 frames, and the nominal defocus range was -0.7 to -1.8 μm. The microscope was optically aligned for fringe-free illumination (Konings et al., 2019), enabling to reduce beam diameter to 0.7 μm on the sample. Imaging was done using an automated low dose procedure implemented in SerialEM (Mastronarde, 2005), in which image shift was used to collect multiple images within a single hole. The beam tilt was adjusted to achieve coma-free alignment when applying image shift.

### Cryo-EM image processing and model building

The GudB_6_-GltAB_2_ Coulomb potential density map was reconstructed in cisTEM (1.0.0-beta) using dose-weighted micrographs. These were prepared by binning the image stacks by a factor of 2, correcting for beam-induced motion, and dose-weighting with MotionCor2 (1.3.0). CTF estimation was performed using CTFFIND4 as part of the cisTEM suite. Micrographs with crystalline ice and poor CTF fits were discarded. Particles were picked in cisTEM using a featureless blob as a template, then were extracted and classified in 2D. All non-ice classes were carried forward to a two-class 3D auto refinement, using an initial reference produced by a cryoSPARC *ab initio* and the default starting resolution of 20 Å. The auto refinement yielded one “noise” class containing poorly aligned particles and one class containing the GudB_6_-GltAB_2_ complex. Particles from this class were carried forward in a single-class 3D auto refinement, followed by manual refinement with C2 symmetry and per-particle CTF correction. To prevent over-refinement, a final high-resolution limit of 3.14 Å was used. The final cisTEM map was auto-sharpened in PHENIX (1.19).

Image processing of the GudB_6_-GltA_6_B_6_ data set was performed using CryoSPARC (3.1.0) (Punjani et al., 2017). The processing scheme is outlined in **Figure S4**. Movies were subjected to patch motion correction, followed by patch CTF estimation. Micrographs with crystalline ice or poor CTF fits were discarded. Particles were initially picked manually from a subset of micrographs. Extracted particles were iteratively classified in 2D and their class averages used as templates for automated particle picking from all selected micrographs. The latter procedure was repeated twice. Particles from well-resolved 2D classes were used for *ab initio* 3D reconstruction and classification, yielding four “noise” 3D classes and one class containing the GudB_6_-GltA_6_B_6_ complex. Particles from this class were carried forward in a 3D non-uniform refinement with imposed D3 symmetry, followed by an additional round of 3D classification to further exclude “noise” particles. The data set was then subjected to local motion correction, per-particle defocus estimation and non-uniform refinement with D3 symmetry. In order to account for movements of asymmetric units within the complex that break the D3 symmetry, we used the ”symmetry expansion” job in CryoSPARC, in which asymmetric units are treated as single particles and rotated in 2D to one of the symmetry-related positions. The newly generated data set contains the asymmetric units of all the complexes and is therefore six times larger. Symmetry expansion was followed by 3D local refinement of the newly generated data set with a binary mask imposed on a single GudB1-GltA1B1 unit and with C1 symmetry. The final map was sharpened in CryoSPARC with a B-factor of -127 before atomic model building.

The initial model for GudB was extracted from PDB 3K8Z, but modified to re-introduce the Q71 residue which appears to be missing in GudB’s published structure. Initial models for GltA and GltB were prepared by homology modeling (SWISS-MODEL) using PDB 1OFD, chain A, as the template for GltA and PDB 6S6T, chain G, as the template for GltB. These models were aligned with the Coulomb potential density map for GudB_6_-GltA_2_B_2_ using rigid body docking (UCSF Chimera 1.14) then refined by performing iterative rounds of manipulation in ISOLDE (ChimeraX 1.1.1, ISOLDE 1.1.0), coot (0.9.3), and phenix.real_space_refine (1.19). The final round of refinement was with phenix.real_space_refine (1.19). From the GudB_6_-GltA_2_B_2_ model, one set of monomers was extracted and refined into the C1 map for the GudB_6_-GltA_6_B_6_ dataset described in the previous paragraph. Refinement similarly involved a round of ISOLDE (ChimeraX 1.1.1, ISOLDE 1.1.0), coot (0.9.3), and phenix.real_space_refine (1.19). To produce the biological assembly, the GudB1-GltA1B1 model was expanded into the full D3 map by applying non-crystallographic symmetry and rigid body docking. The models were inspected in coot (0.9.4.1), ChimeraX (1.1.1) and pyMol (2.4.1). 3D visualization was performed using UCSF Chimera (1.14).

## Supplemental Information titles and legends

**Figure S1.**
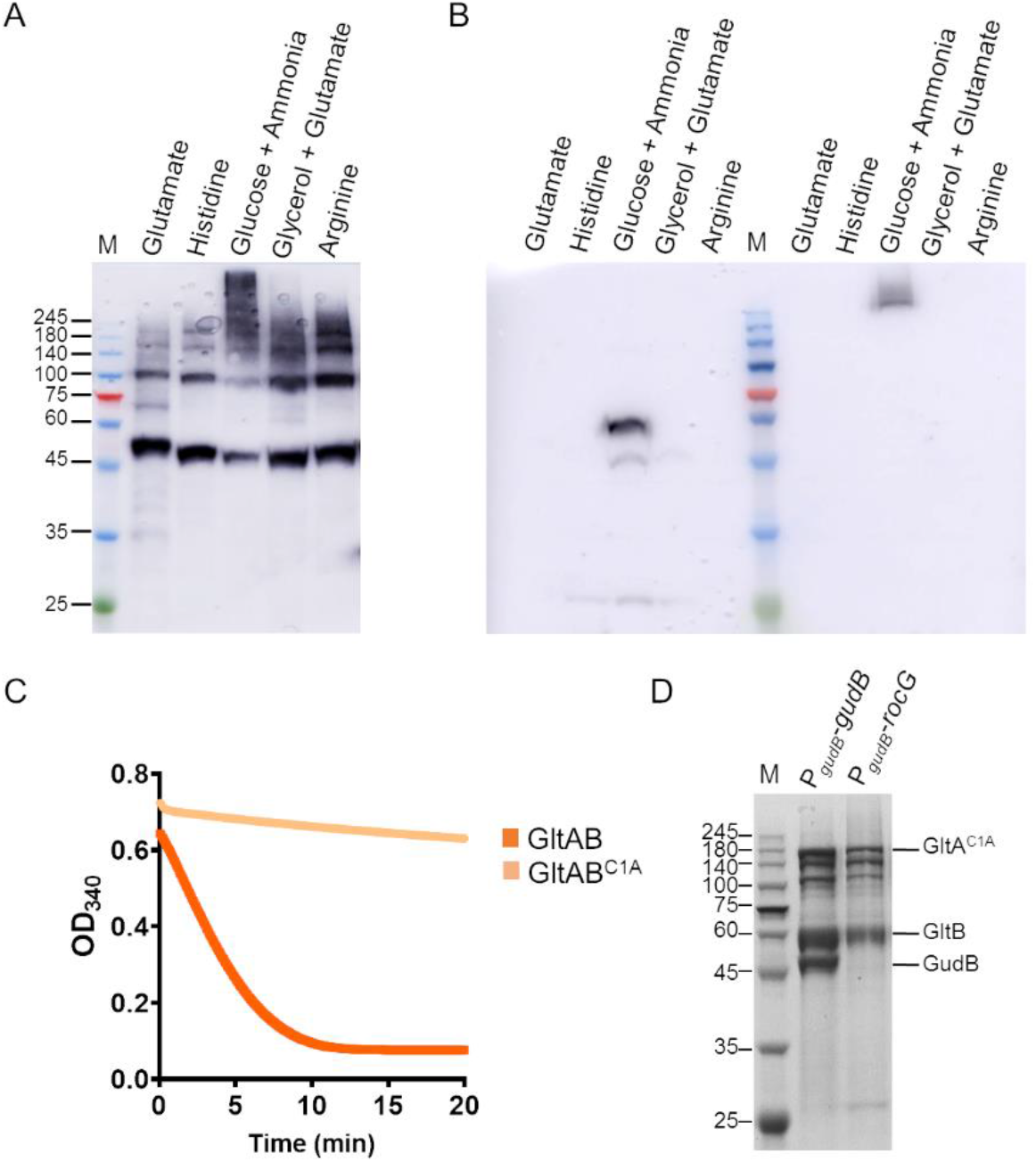
GudB interacts with GltAB. Related to **Figure 1**. **(A)** Western blot analysis using an anti-GudB antibody on lysates prepared from DSG treated *B. subtilis* cells grown on different C/N source. The high molecular weight species of GudB are seen only in cells grown in glucose-ammonia. **(B)** Western blot analysis using a Strep-Tactin-HRP antibody on the same samples used in A. Here the antibody was used on lysates prepared from both DSG-treated (lanes right to the molecular weight marker) and untreated cells (lanes left to the marker). As, can been seen, GltB is expressed only in cells grown on glucose-ammonia, and under this condition, it is also part of a high-MW complex (in the DSG treated sample). Degradation bands of GltB are also seen in the blot and this could contribute to the difference in mass seen in the native-MS (**Figure 4B**) **(C)** Enzyme kinetics of wild-type GltAB (orange) and its mutant GltAB^C1A^ (light orange), show that the mutant is completely inactive. A minor decrease in absorbance with mutant GltA is due to non-enzymatic oxidation of NADPH. The reaction mixture consisted of 2 mM AKG, 5 mM glutamine, 200 µM NADPH, 5 mM DTT, 5 mM MgSO_4_. The reaction was initiated with 2.5 µg of the wild-type or mutant enzyme. **(D)** SDS-PAGE of the eluate from Strep-Tactin column to which lysate from *B. subtilis* strain expressing GltAB^C1A^ in the background of constitutively expressed GudB or RocG is applied. Co-elution of GudB along with the inactive GltA (GltAB^C1A^) upon pulldown of Strep-GltB (P_*gudB*_-*gudB* lane), while GltAB^C1A^ elutes on its own when expressed with RocG (P_*gudB*_-*rocG* lane).

**Table S1.**
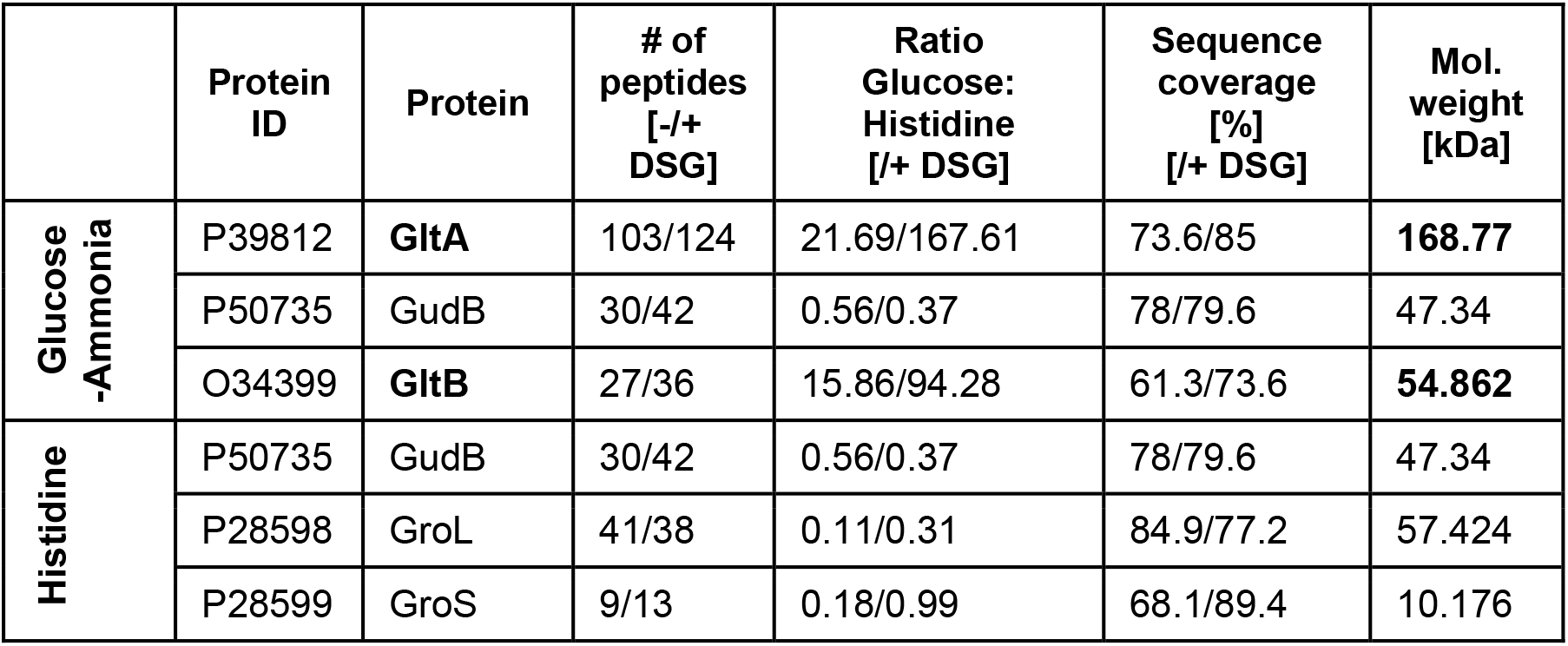
Mass spectrometric identification of GltAB as the interaction partner of GudB. Top hits in the proteomics analysis of the eluates from the immunoprecipitation of GudB from *B. subtilis* cell lysate grown in glucose-ammonia and in histidine, and in the presence and absence of the crosslinker DSG. Both GltA and GltB are highly represented in the glucose-ammonia sample compared to histidine, both in untreated cells and in cells treated with crosslinker (-/+ DSG; the results are shown in this order, *e.g.*, 103 identified peptides without DSG and 124 identified peptides with DSG).

|  | Protein ID | Protein | # of peptides [-/+ DSG] | Ratio Glucose: Histidine [-/+ DSG] | Sequence coverage [%] [-/+ DSG] | Mol. weight [kDa] |
| --- | --- | --- | --- | --- | --- | --- |
| Glucose-Ammonia | P39812 | **GltA** | 103/124 | 21.69/167.61 | 73.6/85 | **168.77** |
|  | P50735 | GudB | 30/42 | 0.56/0.37 | 78/79.6 | 47.34 |
|  | O34399 | **GltB** | 27/36 | 15.86/94.28 | 61.3/73.6 | **54.862** |
| Histidine | P50735 | GudB | 30/42 | 0.56/0.37 | 78/79.6 | 47.34 |
|  | P28598 | GroL | 41/38 | 0.11/0.31 | 84.9/77.2 | 57.424 |
|  | P28599 | GroS | 9/13 | 0.18/0.99 | 68.1/89.4 | 10.176 |

**Figure S2.**
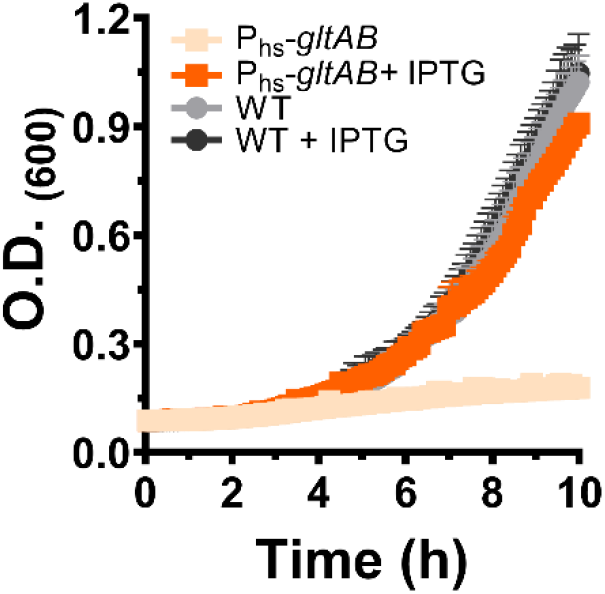
Phenotyping of P_hs_-*gltAB* strain: The strain expresses GltAB from the IPTG-inducible hyperspank (P_hs_) promoter. Without IPTG, the strain cannot grow in minimal medium containing glucose-ammonia as the C/N source (light orange); however, addition of 500 µM IPTG restores growth to almost wild-type levels (orange). Addition of IPTG does not have any effect on the growth of the parental wild-type strain. Error bars indicate standard deviation of three independent measurements.

**Figure S3.**
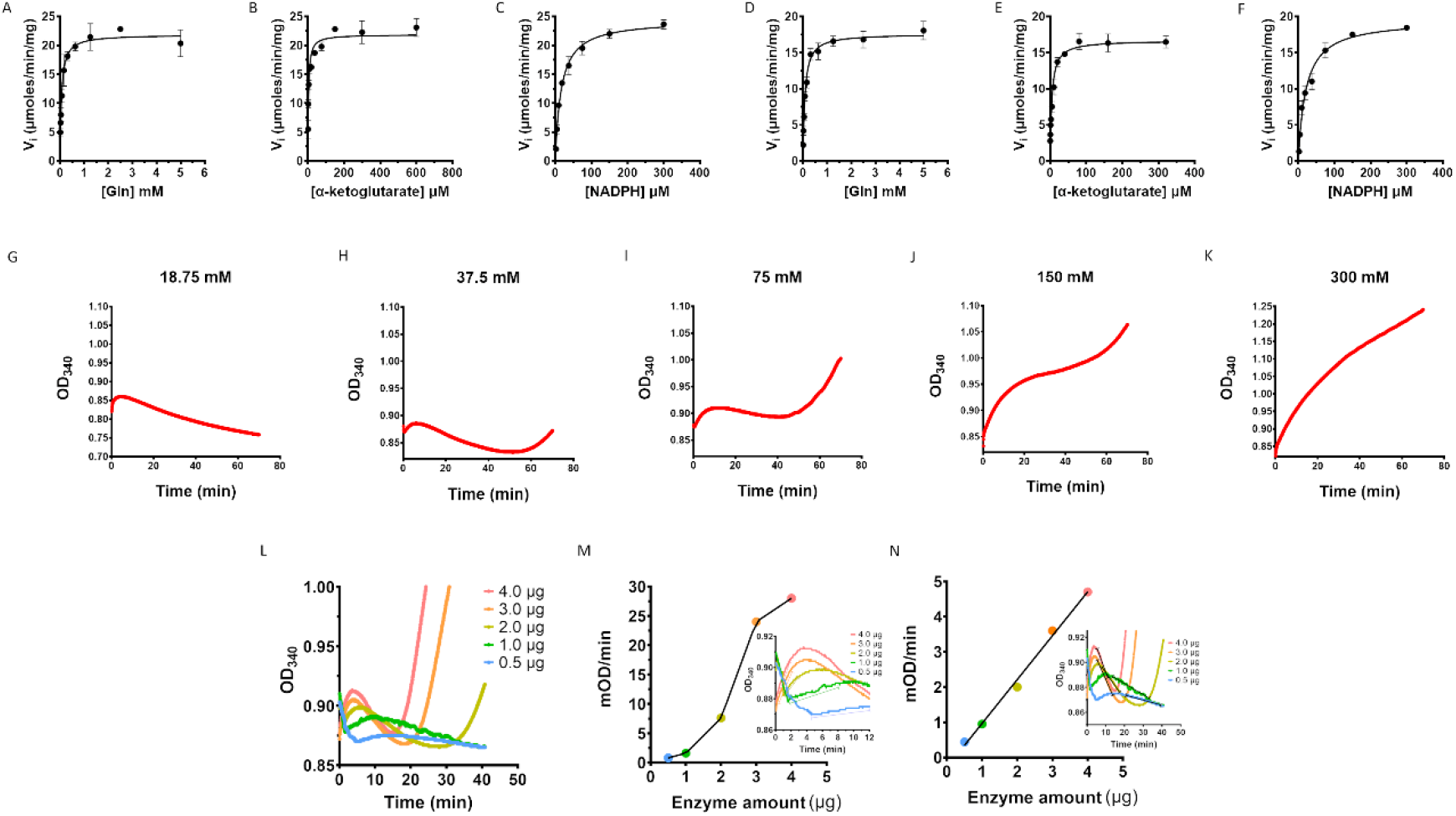
Steady-state kinetics of stand-alone GltAB and of the GudB-GltAB complex. Related to **Figure 3**. **(A-F)** Substrate titration plots for the glutamate synthase activity of GltAB by itself (A-C) and in the GudB bound form (D-F). The activity was monitored by the decrease in absorbance at 340 nm as a result of NADPH oxidation. The assay was performed in degassed 50 mM HEPES buffer pH 7.9 containing 5 mM of magnesium sulfate and 5 mM of DTT. One of the substrates was titrated while keeping the rest saturating: glutamine (5 mM), AKG (α-ketoglutarate, 2 mM), NADPH (200 µM). **(G-K)** Non-hyperbolic progress curves were observed when GudB-GltAB complex activity was monitored in the absence of extraneously added AKG (while all other substrates of GudB and GltAB were present) and at different glutamate concentrations (as denoted above the graphs). GltAB activity is thus now completely dependent on the supply of AKG by GudB’s dehydrogenase activity. While at high glutamate concentrations GudB’s activity dominates (K), at low glutamate concentration the synthase activity takes over (G). At intermediate concentration of glutamate (H-J) the progress curve oscillates between GudB- and GltAB-predominant phases. **(L)** Progress curves obtained using different amount of the GudB-GltAB complex at 37.5 mM glutamate (where oscillations are most pronounced; **Figure S2**, H). The reaction mixture contained all substrates for GudB and GltAB except AKG. M. **(M)** Plotted are the slopes derived from phase 2 of the progress curves (GudB’s activity as shown in the inset) as a function of the complex concentration. The non-linear relationship suggest that association-dissociation of the complex plays a role in turning off-on GudB’s activity. Note also the elapsed time for phase 2 (the 2^nd^ GudB activity phase) that becomes longer as the complex concentration decreases. The inset shows the phase 3 in each of the individual progress curves. **(N)** Unlike phase 2, phase 3 that corresponds to GltAB’s activity shows linear dependence with enzyme concentration. The inset shows phase 3 in each of the individual progress curves.

**Figure S4.**
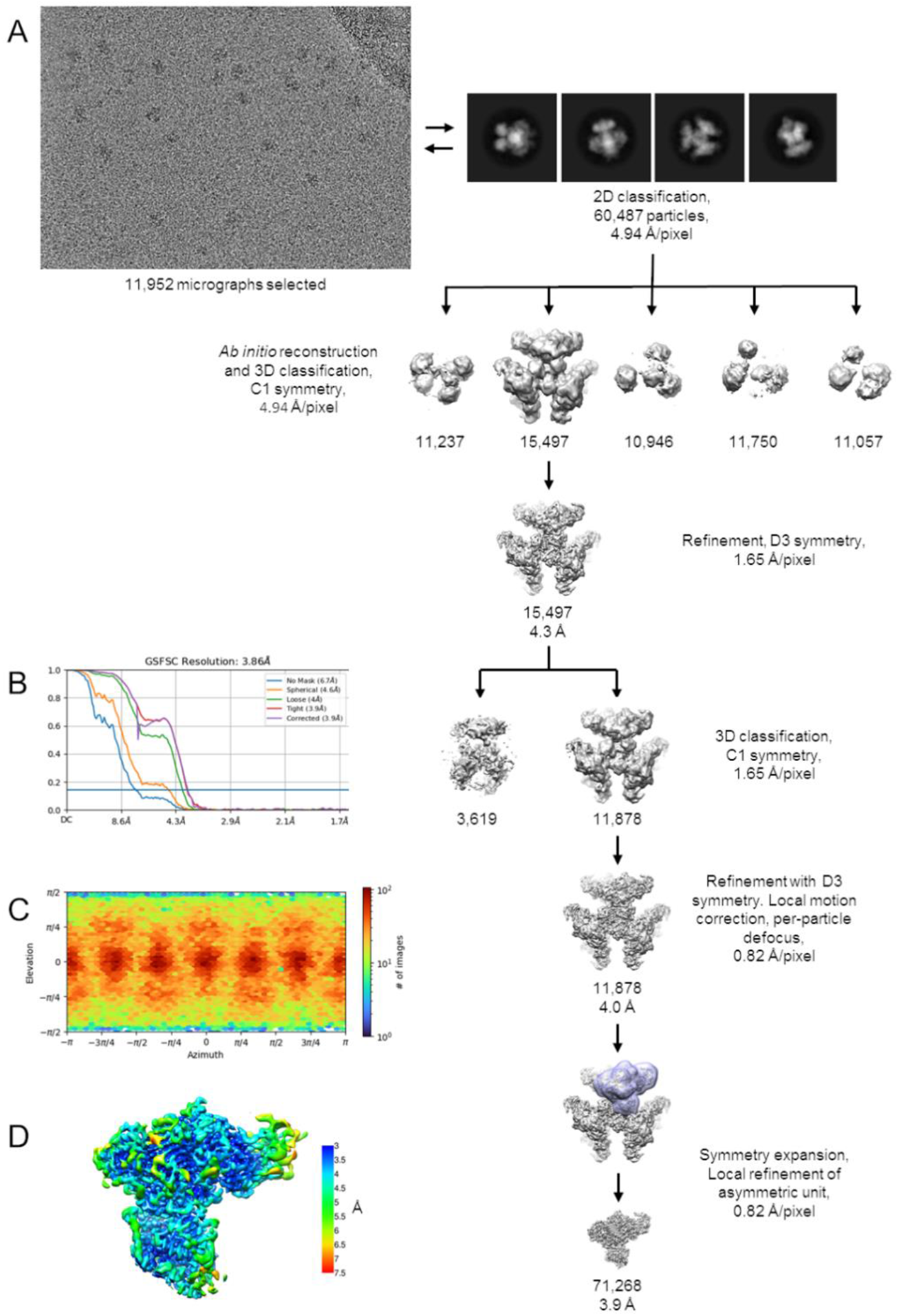
Cyro-EM image processing for GudB_6_-GltA_6_B_6_. **(A)** Scheme for single particle cryo-EM analysis of the GudB_6_-GltA_6_B_6_. Details of the process are described in the Methods section. Briefly, particles were iteratively picked from selected micrographs using well resolved 2D class averages, followed by Ab initio 3D reconstruction and classification into five classes. The best resolved 3D class was refined with D3 symmetry imposed. In order to account for deviations from D3 symmetry, further refinement focused on single GudB-GltAB asymmetric units. The number of particles that are included in the maps are indicated, along with the estimated resolution where relevant. **(B)** FSC curves for the refined GudB-GltAB asymmetric map. **(C)** Angular distribution plot. **(D)** 3D map colored according to local resolution estimate.

**Figure S5.**
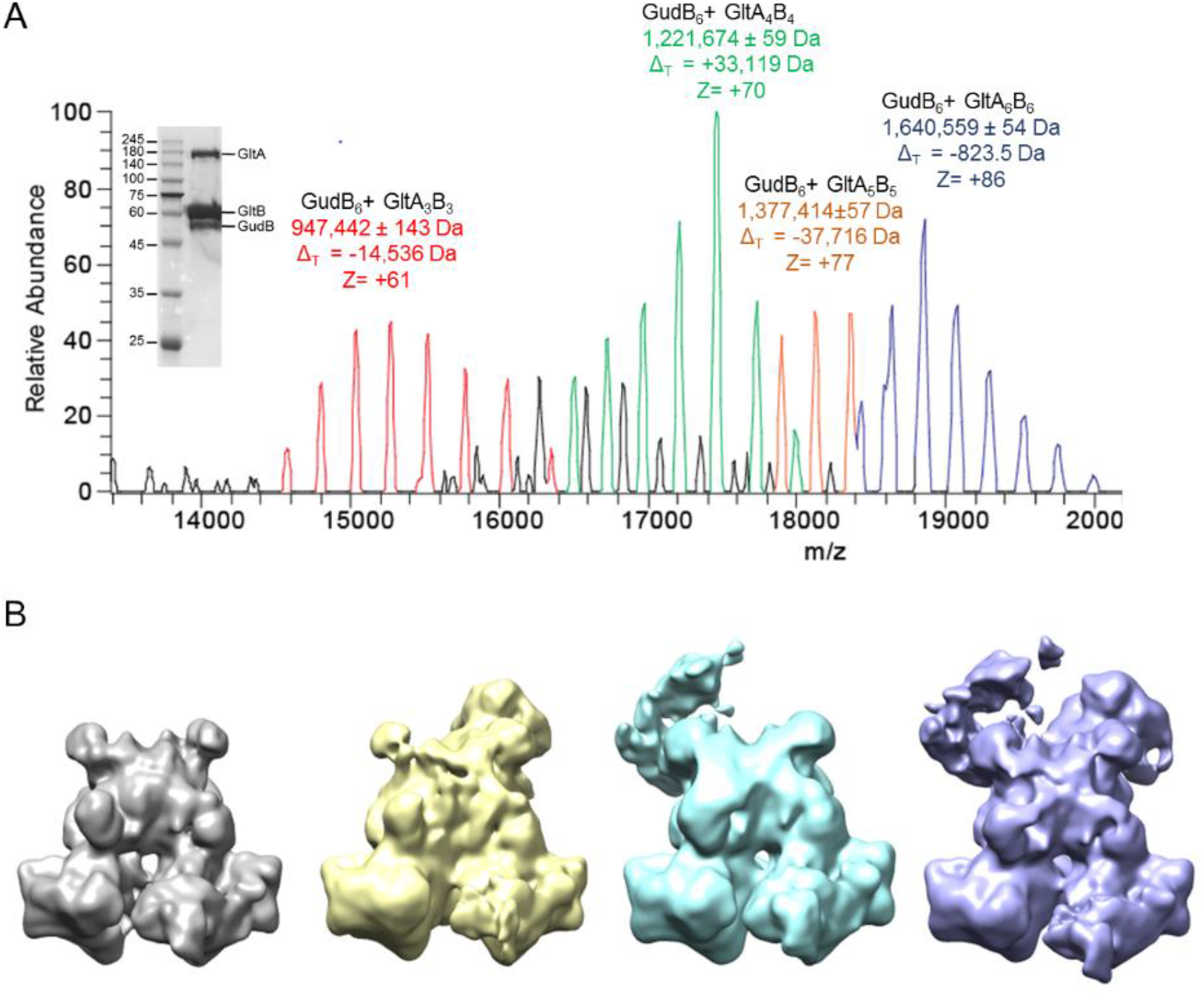
Native-MS and the corresponding particle types observed in cryo-EM of the GudB enriched preparation of the GudB-GltAB complex. **(A)** Native-MS spectra showing different species of GudB-GltAB complex. These species primarily differ in the number of GltAB heterodimers attached to the GudB hexamer (from 3-6). Charge states (z) and the difference from theoretical mass (Δ_T_) is indicated for each species. While the mass of the fully assembled complex agreed well with the expected mass, high Δ_T_ values of the other species could be because of degradation of some of the component proteins during the extended incubation step with *E. coli* lysate (Methods section) during the purification process. The inset shows the SDS-PAGE of GudB-GltAB complex used for the native-MS. The sample was prepared after enriching the *B. subtilis* lysate with recombinant GudB expressed and purified from *E. coli* (see Protein expression and purification, Methods). This sample contained a higher fraction of GudB and was also used for cryo-EM to obtain the high-resolution structure of GudB_6_-GltA_2_B_2_ complex (**Figure 5**). **(B)** Preliminary cryo-EM maps corresponding to different particle types (GudB_6_-GltA_2_-4B_2-4_) observed. The key difference between particles is the number of GltAB subunits – the least being two (grey) and the maximum four (violet).

**Figure S6.**
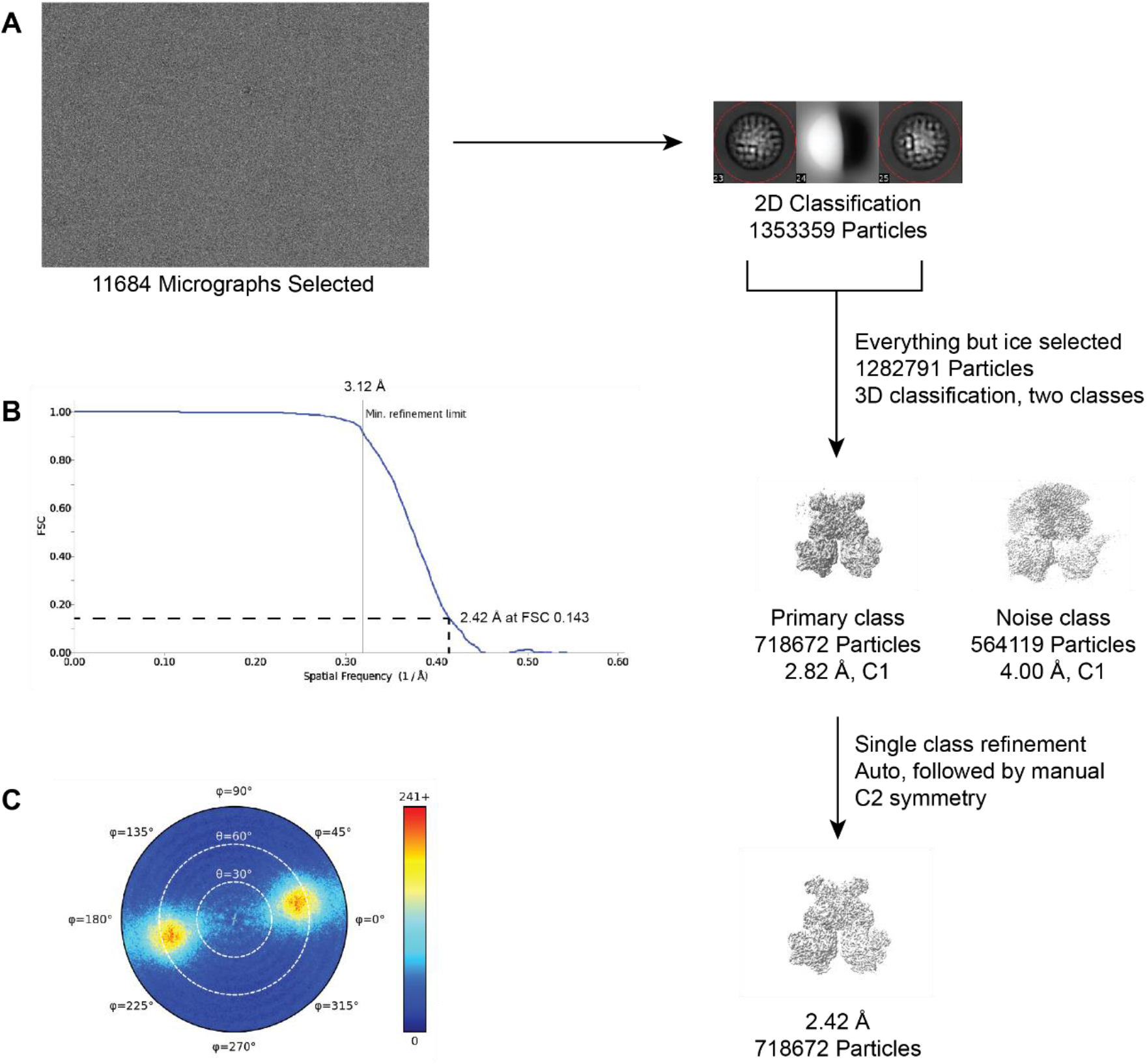
Single particle cryo-EM analysis of the GudB_6_-GltA_2_B_2_. **(A)** The particles were iteratively picked from selected micrographs and classified. All non-ice particles were carried forward and subjected to a two-class 3D auto refinement in cisTEM using the preliminary GudB_6_-GltA_2_B_2_ reference. This yielded one noise class carrying unaligned particles and high frequency noise, and one class representing clear density for the GudB_6_-GltA_2_B_2_ species. This class was refined using auto and manual methods in cisTEM, with C2 symmetry applied. The number of particles and the estimated resolution are indicated **(B)** FSC curves for the refined GudB_6_-GltA_2_B_2_ map. **(C)** Angular distribution plot.

**Table S2.**
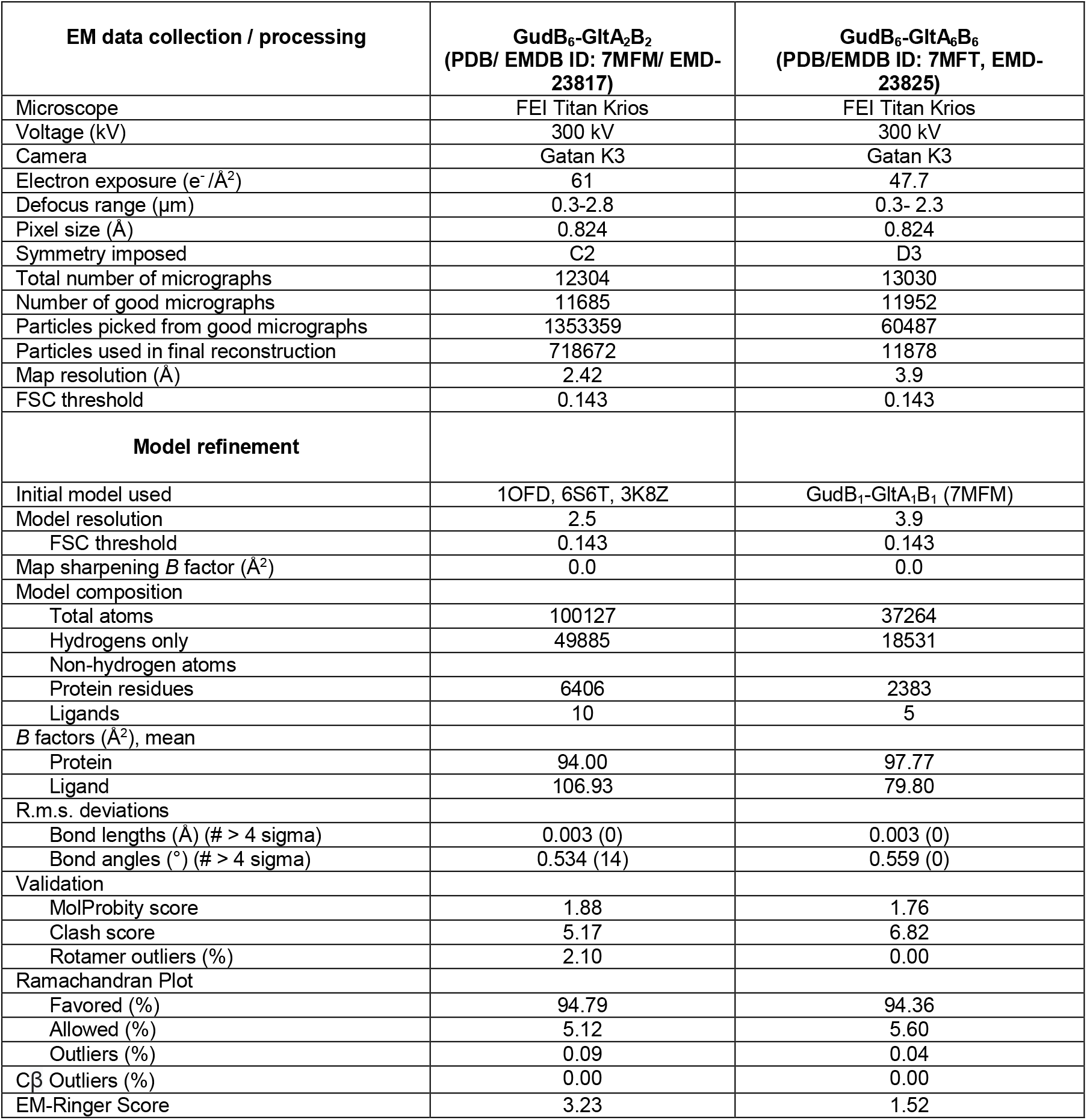
Cryo-EM data collection, processing, and model refinement statistics.

| EM data collection / processing | GudB<sub>6</sub>-GltA<sub>2</sub>B<sub>2</sub><br>(PDB/ EMDB ID: 7MFM/ EMD-23817) | GudB<sub>6</sub>-GltA<sub>6</sub>B<sub>6</sub><br>(PDB/EMDB ID: 7MFT, EMD-23825) |
| --- | --- | --- |
| Microscope | FEI Titan Krios | FEI Titan Krios |
| Voltage (kV) | 300 kV | 300 kV |
| Camera | Gatan K3 | Gatan K3 |
| Electron exposure (e<sup>-</sup>/Å<sup>2</sup>) | 61 | 47.7 |
| Defocus range (μm) | 0.3-2.8 | 0.3- 2.3 |
| Pixel size (Å) | 0.824 | 0.824 |
| Symmetry imposed | C2 | D3 |
| Total number of micrographs | 12304 | 13030 |
| Number of good micrographs | 11685 | 11952 |
| Particles picked from good micrographs | 1353359 | 60487 |
| Particles used in final reconstruction | 718672 | 11878 |
| Map resolution (Å) | 2.42 | 3.9 |
| FSC threshold | 0.143 | 0.143 |
| **Model refinement** |  |  |
| Initial model used | 1OFD, 6S6T, 3K8Z | GudB<sub>1</sub>-GltA<sub>1</sub>B<sub>1</sub> (7MFM) |
| Model resolution | 2.5 | 3.9 |
| FSC threshold | 0.143 | 0.143 |
| Map sharpening *B* factor (Å<sup>2</sup>) | 0.0 | 0.0 |
| Model composition |  |  |
| Total atoms | 100127 | 37264 |
| Hydrogens only | 49885 | 18531 |
| Non-hydrogen atoms |  |  |
| Protein residues | 6406 | 2383 |
| Ligands | 10 | 5 |
| *B* factors (Å<sup>2</sup>), mean |  |  |
| Protein | 94.00 | 97.77 |
| Ligand | 106.93 | 79.80 |
| R.m.s. deviations |  |  |
| Bond lengths (Å) (# > 4 sigma) | 0.003 (0) | 0.003 (0) |
| Bond angles (°) (# > 4 sigma) | 0.534 (14) | 0.559 (0) |
| Validation |  |  |
| MolProbity score | 1.88 | 1.76 |
| Clash score | 5.17 | 6.82 |
| Rotamer outliers (%) | 2.10 | 0.00 |
| Ramachandran Plot |  |  |
| Favored (%) | 94.79 | 94.36 |
| Allowed (%) | 5.12 | 5.60 |
| Outliers (%) | 0.09 | 0.04 |
| Cβ Outliers (%) | 0.00 | 0.00 |
| EM-Ringer Score | 3.23 | 1.52 |

**Table S3.** List of potential interactions between GltA and GudB. Noted are the PDB codes and chain identifiers for which the residue number relate. ^#^ Residues that are different between RocG and GudB. * Residues on the “minor” GudB subunit that interacts less extensively with GltA. H-bond, hydrogen bond; SB, salt bridge.

| **RocG**<br>**(3k92A)** | **GudB**<br>**7MFMA,\*7MFMB** | **GltA**<br>**(7MFMG)** | **Bond type** | **Distance**<br>**(Å)** |
| --- | --- | --- | --- | --- |
| R124 | R124 | K573 | H-bond | 2.94 |
| N231 | N231 | N447 | H-bond | 2.77 |
| D270 | D270 | K330 | SB | 3.06 |
| R272 | R272 | Q445 | H-bond | 2.41 |
| D273 | D273 | Y560 | H-bond | 3.34 |
| S274 | S274 | K558 | H-bond | 3.30 |
| N280<sup>#</sup> | **K280** | K708,E324 | H-bond, SB | 3.01,3.08 |
| R84 | R84 | E451 | SB | 3.45 |
| H86 | H86 | E454 | SB | 3.66 |
| K271<sup>#</sup> | **R271** | E324 | SB | 3.00 |
| H398<sup>#</sup> | **\*R398** | E551,<br>S550 | SB, | 3.66 |
| K399<sup>#</sup> | **\*R399** | Y433 | H-bond/Cation-Pi | 2.97 |
| R419 | **\*R419** | D579 | SB | 2.56 |

**Figure S7.**
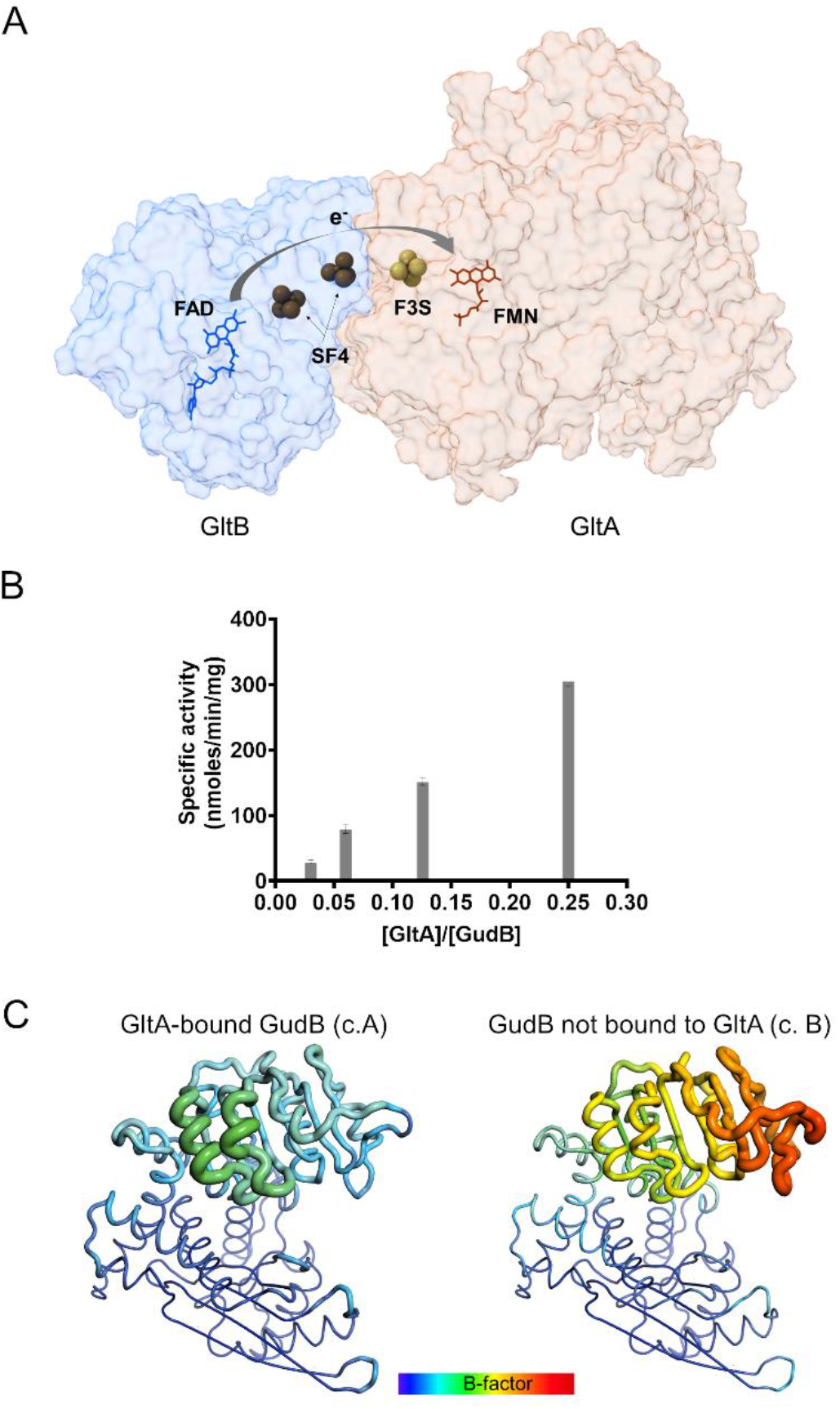
Details of the GudB-GltAB interaction. **(A)** The co-factors in GltAB: FAD, two 4Fe-4S clusters (SF4), 3Fe-4S cluster (F3S) and FMN. These co-factors are involved in shuttling of electrons from NADPH to 2-iminoglutarate along shown arrow. GltA and GltB are shown in a transparent surface representation. **(B)** GltAB in sub-stoichiometric amounts promotes hexamerization of GudB by interacting with multiple GudB protomers (from different dimers, as shown in **Figure 5D**) and hence prevents loss of activity. The assay buffer contained 400 mM glutamate and 4 mM NAD^+^ and the reaction was initiated by the addition of an enzyme mix consisting of GudB pre-incubated with different amounts of GltAB (in all the reactions, GudB was present at a final concentration is 0.05 µM). GltAB prevents GudB inactivation in a concentration dependent manner. Error bars indicate standard deviation of two independent measurements. **(C)** Binding to GltA stabilizes many loops in the cofactor binding domain of GudB. Shown are a GltA-bound GudB protomer (left, chain A in 7MFM) and a free GudB protomer in the same structure (right, chain B) in the “putty” representation as implemented in PyMol. The radius of the ribbon increases from low to high B-factor and the Cα B-factors are shown in dark blue (lowest B-factor, 54) to red (highest B-factor, 163).

**Figure S8.**
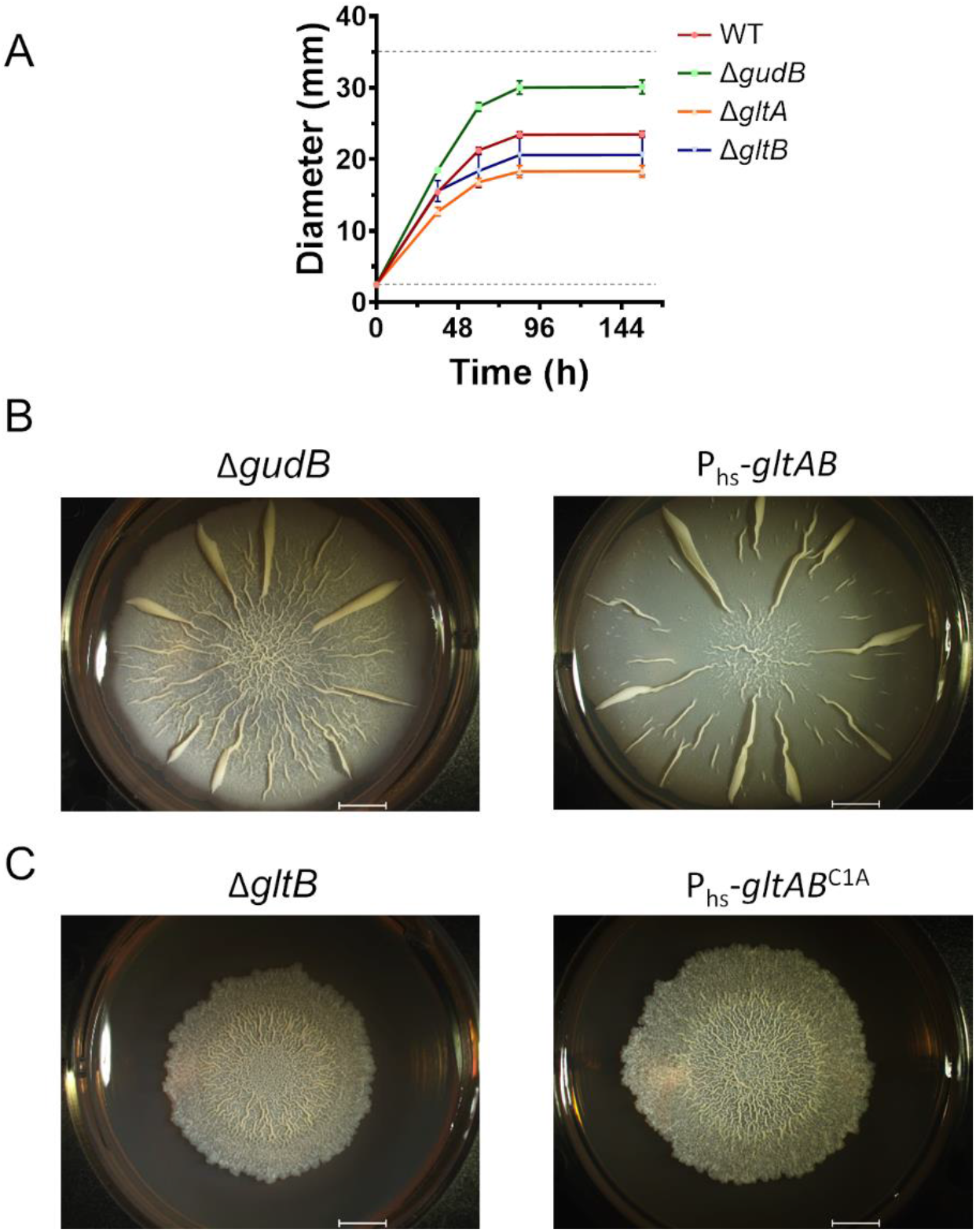
Biofilm growth and disruption phenotypes. **(A)** Biofilm diameters measured at different time points. While ΔgudB biofilms were bigger in size and grew faster than wild-type biofilms, Δ*gltA* and Δ*gltB* biofilms were smaller. The dashed lines at 2.5 mm and 35 mm indicates the starting size of the biofilm, and the diameter of the well used to grow the biofilm, respectively. Error bars represent standard deviation of 4 independent measurements. **(B)** Similarity in biofilm morphology between Δ*gudB* and the P_hs_-*gltAB* strains (grown with 100 µM IPTG). Overexpression of GltAB in the latter increases in synthase activity and also silences GudB, thereby resembling the Δ*gudB* biofilm morphology. Both biofilms grew rapidly and had large wrinkles spreading from the interior to the periphery of the biofilm. **(C)** Similarity in biofilm morphology between Δ*gltB* and the P_hs_-*gltAB*^C1A^ strains (grown with 100 µM IPTG). In both the biofilms the wrinkles are restricted to the interior of the biofilm.

**Appendix Table A1.** *B. subtilis* strains used in the study

| Strain name | Genotype | Description |
| --- | --- | --- |
| 3610 | Prototroph | Wild-type *Bacillus subtilis* |
| *gltB*-TS | 3610 *gltB::gltB-TS* | GltB, under its native promoter, tagged with a twinstrep (TS)-at its C-terminus. Used for pulldown of GltB and of the GudB-GltAB complex for kinetic studies. |
| *gltB*-TS-$P_{gudB}^-$ *rocG* | 3610 *gltB::gltB-TS rocG::kan amyE::(P<sub>gudB</sub><sup>-</sup>-rocG-spec)* | As above, *plus* RocG placed under GudB's promoter (at the *amyE* landing site). Endogenous *rocG* is knocked out with a kanamycin cassette. Used for testing the in vivo interaction capability of RocG with GltAB |
| Δ*gudB* | 3610 *gudB::erm* | Knockout of *gudB*. Used for growth profiling in media containing different C/N source |
| Δ*gltA* | 3610 *gltA::erm* | Knockout of *gltA*. Used for growth profiling in media containing different C/N source |
| Δ*gltB* | 3610 *gltB::erm* | Knockout of *gltB*. Used for growth profiling in media containing different C/N source |
| Δ*gltABC* | 3610 *gltABC::erm* | A knockout of the entire *gltABC* operon. Used for the generation of all the following strains. |
| $P_{hs}^-$-*gltAB*<sup>C1A</sup> | 3610 *gltABC::(P<sub>hs</sub><sup>-</sup>-gltA<sup>C1A</sup>-gltB-TS-cat)* | Strain expressing the inactive mutant of GltA (GltA<sup>C1A</sup>) along with a GltB-TS from an IPTG inducible hyper spank promoter ($P_{hs}^-$). GudB is expressed from its endogenous promoter ($P_{gudB}$). Used for planktonic and biofilm growth phenotyping and for the purification of the GudB-GltAB<sup>C1A</sup> complex. |
| $P_{hs}^-$-*gltAB*<sup>C1A</sup>+$P_{gudB}^-$ *rocG* | 3610 *gltABC::(P<sub>hs</sub><sup>-</sup>-gltA<sup>C1A</sup>-gltB-TS-cat) rocG::kan amyE::(P<sub>gudB</sub><sup>-</sup>-rocG-spec)* | Strain expressing the inactive mutant of GltA (GltA<sup>C1A</sup>) along with a GltB-TS from $P_{hs}^-$. The strain also constitutively expresses RocG using $P_{gudB}^-$ from *amyE* locus. Used for planktonic and biofilm growth phenotyping and for the purification of the GudB- |
|  |  | GltAB<sup>C1A</sup> complex. The strain was also used to test the *in vivo* interaction capability of RocG with GltAB<sup>C1A</sup>. |
| $P_{hs}$-gltAB | 3610 gltABC ::<br>($P_{hs}$-gltA-gltB-TS) | Strain expressing GltA along with a GltB-TS from $P_{hs}$. Used for overexpression and purification of functional GudB-GltAB complex for structural studies. The strain was also used for planktonic and biofilm growth phenotyping. |
| $P_{hs}$-gltAB+$P_{gudB}$-rocG | 3610 gltABC ::<br>($P_{hs}$-gltA-gltB-TS)<br>rocG::kan amyE ::<br>($P_{gudB}$-rocG-spec) | Strain expressing GltA along with a GltB-TS from $P_{hs}$. The strain also constitutively expresses RocG using $P_{gudB}$ from the amyE locus. The strain was also used for planktonic and biofilm growth phenotyping. |
| $P_{hs}$-gltAB -<br>ΔgudB | 3610 gltABC ::<br>($P_{hs}$-gltA-gltB-TS)<br>gudB::erm | Strain expressing GltA along with GltB-TS from $P_{hs}$ but devoid of GudB. For overexpression and purification of GltAB without co-purification of GudB. |

**Appendix Table S2.**
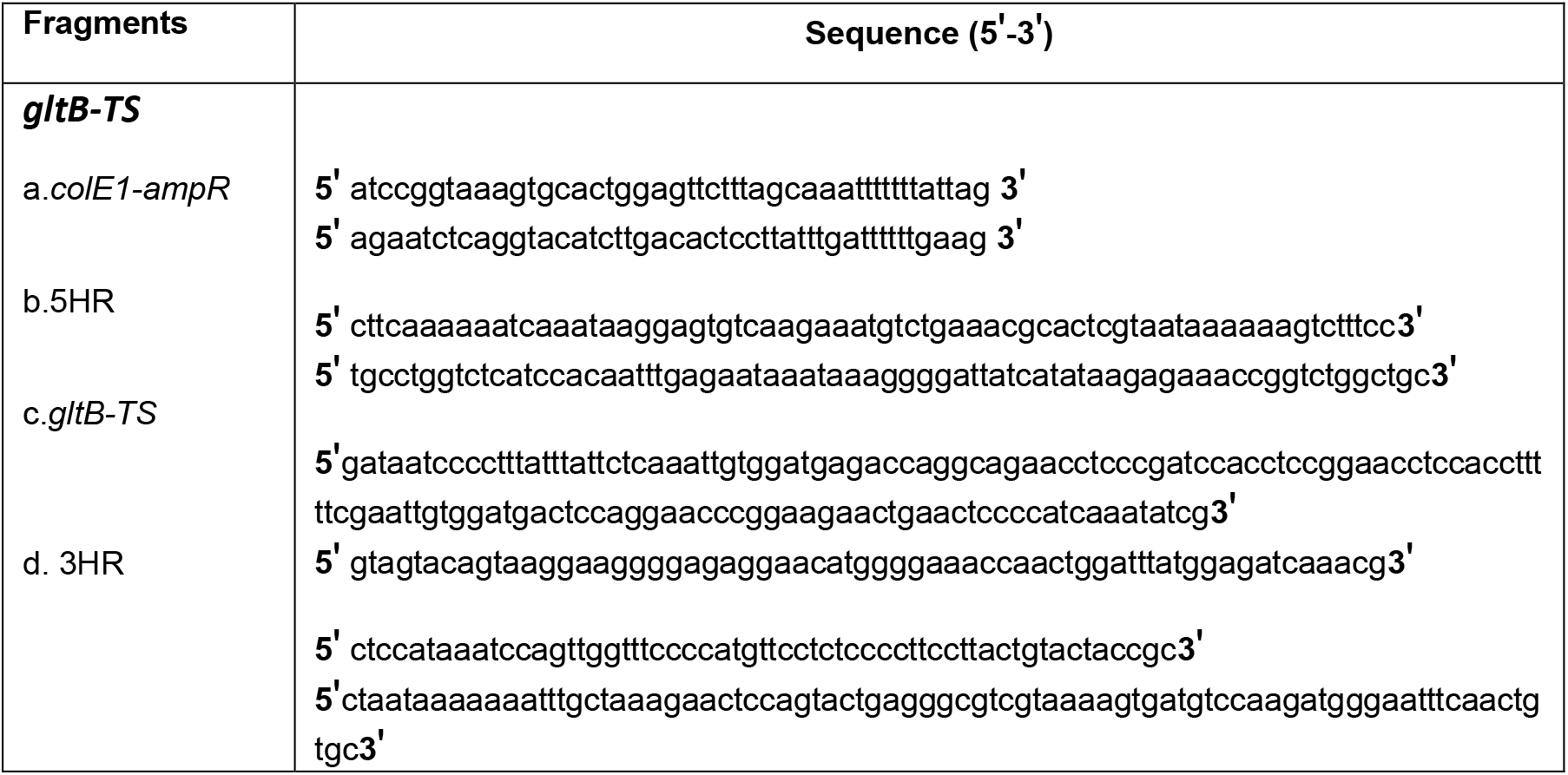
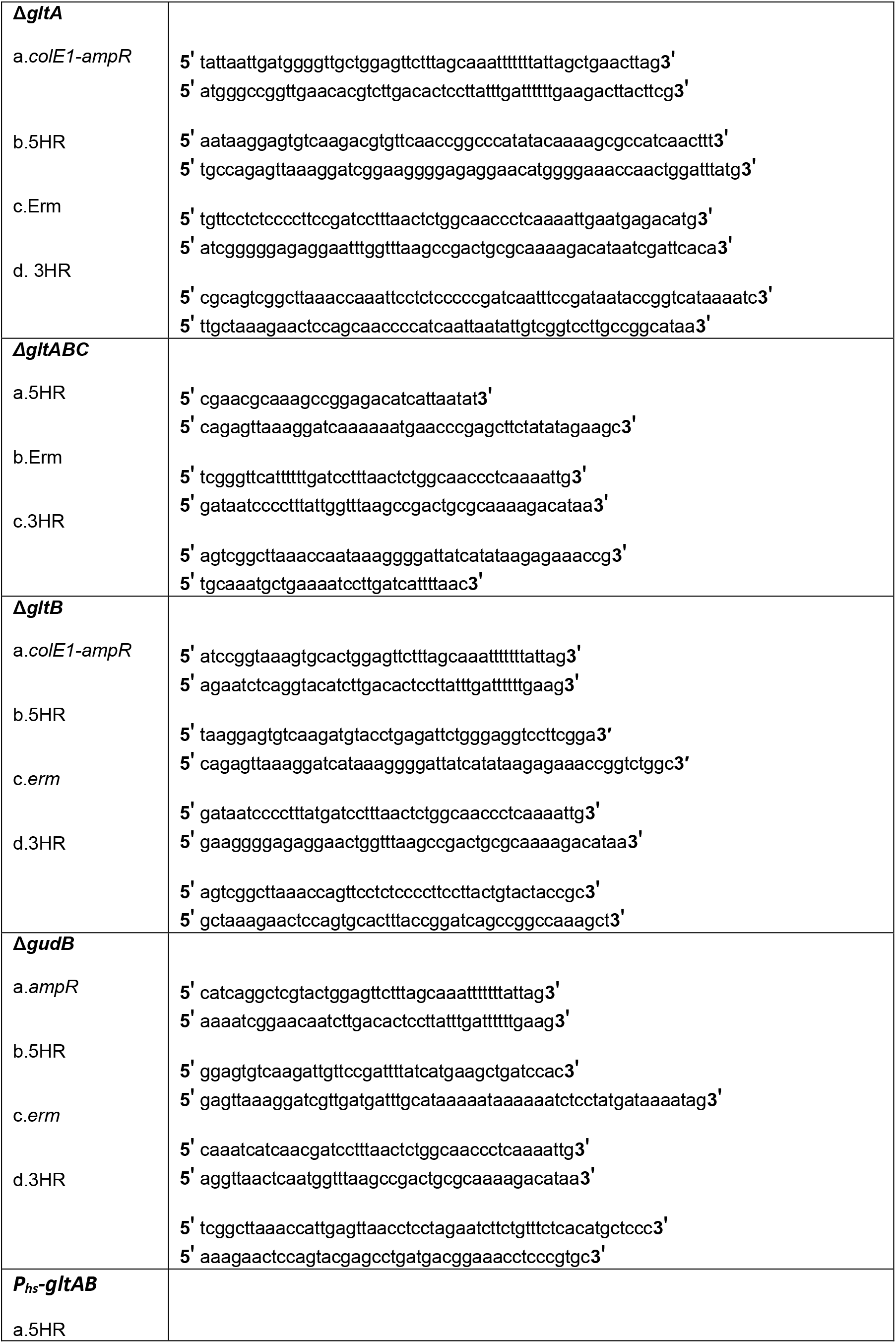
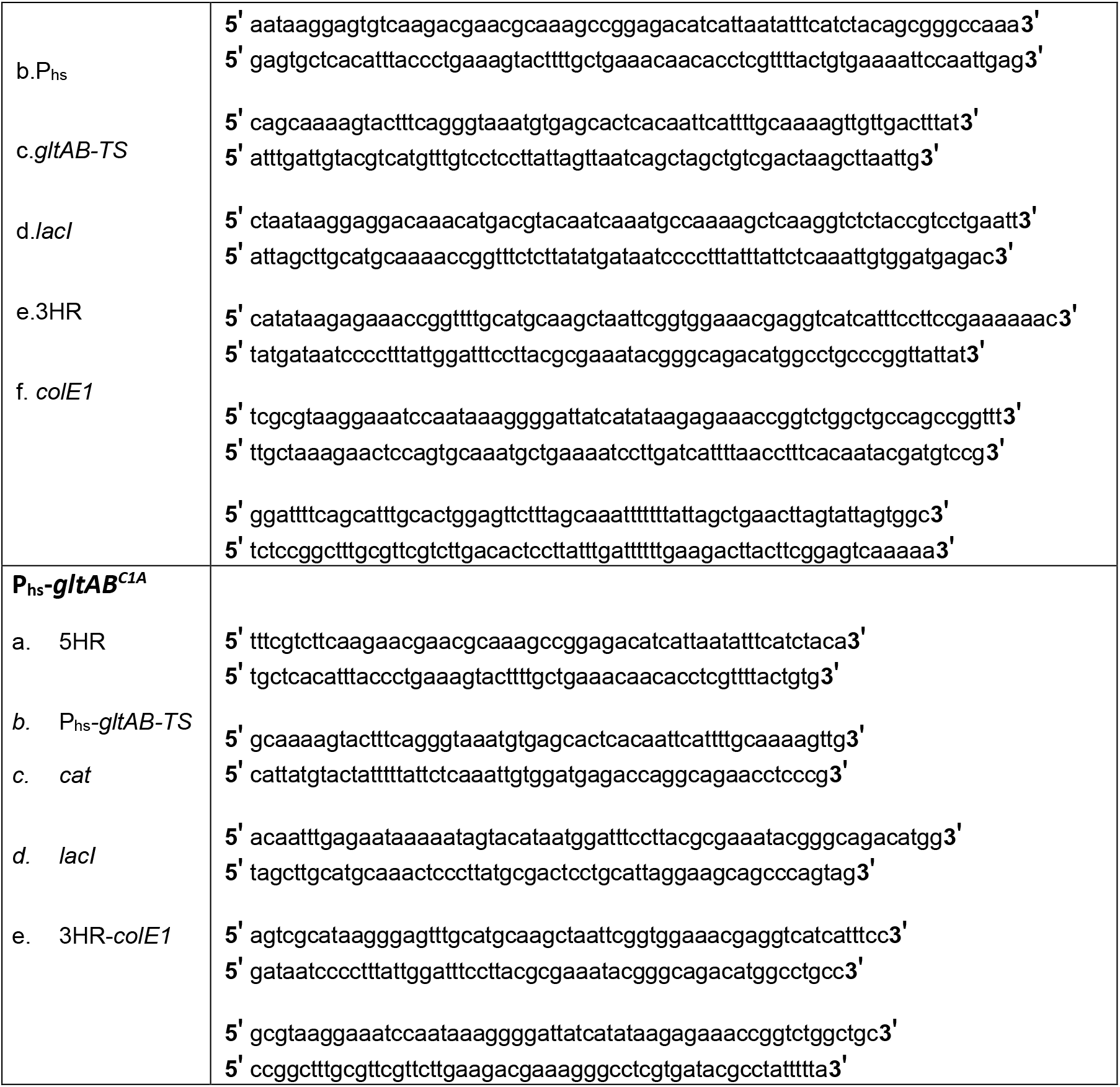
Oligonucleotides used in the study. Each row of the table provides the name of the construct, the fragments used the generate the construct and oligonucleotide sequence used to generate the respective fragment. 5HR and 3HR corresponds to the 5ʹ and 3ʹ homology arms for the recombination.

